# Single cell and spatial characterization of the human pancreas reveals drivers of beta cell dysfunction in cystic fibrosis

**DOI:** 10.64898/2025.12.14.694120

**Authors:** Hannah M. Mummey, Sierra Corban, Jacinta Lucero, Rebecca L. Melton, Madeleine Pittigher, Victoria C. Johnston, Gail H. Deutsch, Agnieszka D’Antonio-Chronowska, Allen Wang, Rebecca L. Hull-Meichle, Kyle J. Gaulton

## Abstract

Cystic fibrosis (CF) causes severely damaged pancreas morphology and results in high rates of cystic fibrosis-related diabetes (CFRD), but the cellular pathways and regulatory programs driving CFRD pathogenesis in the pancreas are not understood. In this study, we performed single cell multiomic and spatial profiling of 2.04M cells in the pancreas from 23 non-disease and CF donors and defined regulatory programs and tissue niches of specific cell types and sub-types in CF. A high-mucin sub-type of ductal cells had relatively preserved abundance, altered localization and up-regulated stress, secretion and pro-fibrotic activity in CF, and conventional ductal cells showed evidence for transition to the high-mucin state. Increased inflammatory and fibrotic pathway activity in CF was linked to closer proximity and crosstalk between immune and stellate cells in specific niches. Beta cells had extensive genomic changes in CF, including increased stress and insulin secretion-related processes, and these changes were broadly distinct from those in type 1 and type 2 diabetes. Islets in CF preferentially localized near large adipose tissue and collagen structures, and both areas were strongly linked to beta cell loss in CF due to signaling from adipocytes, macrophages, stellate, and high-mucin ductal cells. Overall, our results reveal cellular drivers of pancreatic dysfunction in CF and offer new in-roads to preserving beta cells in CFRD.

## Introduction

Cystic fibrosis (CF) is an autosomal recessive disorder caused by mutations of the *CFTR* gene^1–4^. Individuals with CF experience mucus build up in the lungs and are at increased risk of infection from air-borne pathogens, and respiratory failure is the leading cause of death in CF^5^. However, individuals with CF also experience systematic damage to many other tissues including the pancreas, intestine, kidneys, liver, and vas deferens^6–9^. In the past decade, quality of life and lifespan have increased due to the development of small molecule therapies (highly effective CFTR modulators; HEMT) that correct over 90% of known *CFTR* folding and conductance errors ^10,11^. However, in parallel with increases in life expectancy, the prevalence of serious age-related complications involving other tissues has also increased.

Individuals with CF have long been known to have altered pancreatic morphology with substantial exocrine loss and replacement with fibrosis and fatty accumulation^12^, and ∼85% of people living with CF rely on digestive pancreatic enzyme therapy to compensate for the loss of enzyme-producing acinar cells^13^. Cystic fibrosis-related diabetes (CFRD) is linked to this altered tissue morphology although the underlying mechanisms of disease are poorly understood. Studies have estimated that half of adults with CF develop CFRD, with higher rates tied to more severe *CFTR* mutations^14,15^, suggesting that CFRD is a complex disease with incomplete penetrance. Compared to those with CF and normal glucose tolerance and insulin secretion, individuals with CFRD have lowered lung function^16^, higher airway glucose concentrations which can lead to increased growth of respiratory pathogens^17^, a substantial increase in disease and mental health burden^18,19^, and higher mortality rates^15^. Studies of the impact of HEMT on CFRD have thus far been inconclusive^20,21^. A greater understanding of the mechanisms leading to CFRD and the development of treatments to prevent CFRD is therefore critically important.

Reduced beta cell mass has been found in CF islets starting from a young age^22,23^ and individuals with CF have reduced endocrine function^24–26^, although islet structures remain intact and populations of beta cells remain in CF^22,23,27–31^. Genetic risk of CFRD is linked to insulin secretion-related loci involved in type 2 diabetes^32^, suggesting that reduced beta cell function is a risk factor for CFRD. Increased glucagon immunoreactivity has been found in CFRD islets^22,23,31,33^ indicating changes to alpha cells as well as well beta cells, although there is evidence that glucagon secretion is also impaired in CF^23–25^. Pancreatic ductal cells highly express *CFTR* with low expression in other pancreatic cell types^23,34,35^, and deleting *CFTR* in beta cells in mice had no impact on insulin secretion or key beta cell genes^23^. Therefore, the initiating drivers of beta cell dysfunction and CFRD due to *CFTR* mutations are potentially external to beta cells, indicating interactions with other cell types and the tissue environment. Increased immune infiltration and islet amyloid deposition has been found in islets in CFRD^22,23^, implicating pro-inflammatory processes in beta cell dysfunction. However, a comprehensive understanding of the specific changes within beta and other endocrine cells in CF - as well as the external cell types, structures and interactions in the pancreas driving these changes - is lacking.

Single cell and spatial genomics enable profiling the gene regulatory programs, signaling networks and localization of cell types in a heterogeneous tissue sample. There have been few single cell genomics studies performed in CF in the airways, liver, and kidney^36–38^ with none in the pancreas in humans or model organisms, and previous imaging-based studies of the pancreas in CF have had limited resolution and depth^22,23,31,39^. In this study, we generate the first single cell-resolved map of gene expression, gene regulation, and spatial organization in the pancreas during CF using multimodal single nuclear sequencing and imaging-based spatial transcriptomics. We identified changes in abundance, localization and gene regulatory programs in many pancreatic cell types in CF, and identify the specific structures and signaling networks that drive beta cell dysfunction and death in CF.

## Results

### Single cell and spatial profiling reveal large-scale cellular changes in the pancreas in CF

We profiled pancreatic tissue from 23 individuals with cystic fibrosis (CF) (n=10) or age- and sex-matched controls (n=13) using 10x multiome (paired snRNA-seq+snATAC-seq) and Nanostring CosMx assays (**Figure 1a, Supplementary Table 1**). Of the 23 profiled individuals, we performed 10x multiome assays in 19 donors (9 CF, 10 control) and CosMx assays and H&E staining in 12 donors (6 CF, 6 control), where eight donors were profiled with both technologies. The H&E images of 10 donors were further annotated by a board-certified pathologist (**Figure 1b**). Most CF donors were diagnosed with CFRD (n=7); however, a small subset did not have a diabetes diagnosis (CFND) (n=3). In addition, all CF donors had class I or class II *CFTR* mutations, which have the most severe impact on protein expression and function and lead to severe disease pathology^40^, although for one donor (nPOD_CF1) only one of the *CFTR* alleles was known.

**Figure 1.**
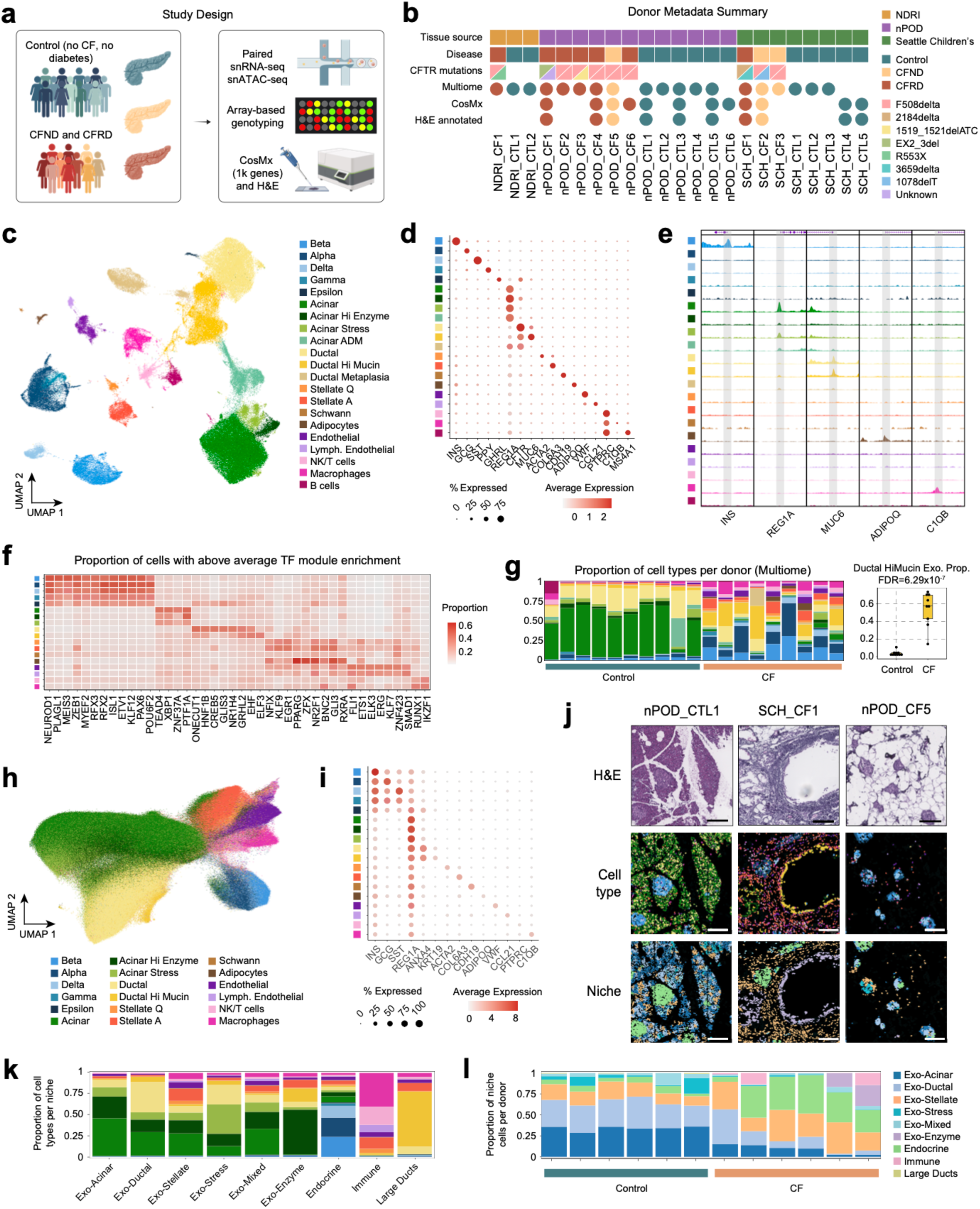
Pancreatic cell type composition and organization changes during cystic fibrosis. a) Illustration of study design. Pancreatic tissue samples were obtained from donors without Cystic Fibrosis or diabetes (control), Cystic Fibrosis but no diabetes (CFND) and Cystic Fibrosis Related Diabetes (CFRD). Samples were profiled using a combination of paired single nuclear RNA-seq and single nuclear ATAC-seq, array-based genotyping and spatial transcriptomics. Created with BioRender. b) Summary of donor metadata including tissue source, disease and mutation status and assays used to profile donor samples. c) UMAP projection of 120,075 10x Multiome barcodes after QC filtering, demultiplexing and doublet removal. A joint embedding was created after performing dimensionality reduction separately on both RNA and ATAC data. d) Per-cell type gene expression of pancreatic marker genes from Multiome map. The dot color represents the average expression in a cell type, and the dot size represents the percentage of cells in the cell type with non-zero expression. e) ATAC signal tracks for select cell type-marker genes. Each window contains a 6kb region centered around the marker gene transcription start site (TSS) and peak heights are standardized across all cell types for each region. f) Heatmap of transcription factor module enrichment summarized across cell types. Per-cell type enrichment was calculated as the proportion of cells per cell type with per-cell enrichment AUC values more than 1 standard deviation above the overall mean. g) Left: proportion of total Multiome cells for each donor classified by CF status (blue=Control, orange=CF) and colored by cell type. Right: within exocrine compartment proportions of high-mucin ductal cells split between control and CF donors. Significance value is based on a t-test. h) UMAP projection of 1,924,573 cells profiled via Nanostring CosMx after QC filtering, colored by cell type identities assigned via scANVI integration and prediction. i) Per-cell type gene expression of pancreatic marker genes from CosMx map. The dot color represents the average expression in a cell type, and the dot size represents the percentage of cells in the cell type with non-zero expression. j) Example images of three donors profiled with CosMx and stained post-CosMx with H&E. Top row: H&E staining; middle row: CosMx segmented cells colored by scANVI predicted cell type; bottom row: CosMx segmented cells colored by BANKSY niche. Scale bars represent 200µm. k) Proportion total cells per BANKSY niche colored by cell type. l) Proportion of total CosMx cells for each donor classified by CF status (blue=Control, orange=CF) and colored by niche.

We performed single cell multiome assays on pooled nuclei representing 18 CF and control donors, where cells from each assay were demultiplexed to donor-of-origin using array genotyping data (**see Methods**). We also performed multiome assays for three donors (1 CF, 2 control) from NDRI individually due to large amounts of available tissue, including one control donor not profiled in the pooled experiments. After barcode filtering with strict quality control metrics (**see Methods**), we clustered 120,075 nuclei (65,304 control, 54,771 CF) using weighted nearest neighbors in Seurat^41^ (**Figure 1c**). The final map consisted of 21 clusters, which were annotated for cell type identity based on the expression and chromatin accessibility of known cell type marker genes (**Figure 1d-e, Supplementary Figure 1a**). In addition to previously reported exocrine, endocrine, immune, endothelial, and stellate populations, due to the dramatic changes in CF pancreas, we identified multiple other cell types and sub-types not seen in previous pancreas maps. For example, we identified adipocytes, marked by expression of *ADIPOQ* as well as markers of mature adipocytes such as *PPARG*, *FABP4*, and *PLIN1,* which were observed only in CF due to extensive adipose accumulation in these individuals (**Figure 1c-g**, **Supplementary Figure 1b, Supplementary Table 2**). We also identified epsilon cells, marked by the expression of *GHRL,* which are a rare population of endocrine cells. Several exocrine sub-types with differentiation-related profiles were comprised of cells from a single CF donor, and thus we did not include them in further analyses (**Supplementary Figure 1c,d**).

We defined the gene regulatory programs of each pancreatic cell type and sub-type identified in our study. We identified *cis*-regulatory elements (cREs) in each cell type and defined a unified set of 228,495 cREs across all cell types. We further defined gene regulatory networks (GRNs) of transcription factor (TF) activity in each cell type by leveraging the paired snRNA-seq and snATAC-seq profiles with SCENIC+^42^ (**Supplementary Data 1**). We identified a total of 275 TF-centered GRNs, henceforth referred to as TF modules, of which 164 were based on motifs with direct evidence for TF binding and 111 on extended evidence. TF modules with increased activity in specific cell types revealed likely regulators of cell type identity and function. We identified 42 cell type-enriched TF modules (**Supplementary Table 3**), which included many factors known to drive cell type-specific regulatory activity, including *NEUROD1* and *RFX* TFs in endocrine cells, *ONECUT1* and *HNF1B* in ductal cells, and *RUNX1* in immune cells (**Figure 1f**). In addition, we identified regulators of cell types and sub-types not extensively described in previous single cell maps of the human pancreas including *PPARG, RXRA* and *ZFX* in adipocytes and *EHF* and *ELF3* in high-mucin ductal cells.

Cellular composition was dramatically altered in the CF pancreas. As expected, we observed a dramatic decrease in acinar cell populations across CF donors, although many other populations were altered as well. For example, there was a substantial increase in the proportion of the high-mucin ductal sub-type (**Figure 1c-e**), which has been described in previous single cell maps of the human pancreas with very low abundance^35,43,44^ relative to the conventional ductal population in CF (**Figure 1g**). Additionally, there was a large increase in pro-fibrotic and inflammatory cell populations in CF (**Figure 1g**). As the increased abundance of many cell types is likely related to the decrease in acinar cells, we confirmed significant changes in proportions of many cell types by re-performing these analyses within compartments. We found a significant decrease in acinar cells compared to all exocrine cells, and significantly increased within-compartment proportions of high-mucin ductal cells, adipocytes, and endothelial cells (**Figure 1g**, **Supplementary Figure 2a-b, Supplementary Table 4**).

Next, we processed and applied quality control filters to spatial transcriptomics data from CF and control individuals, resulting in 1,924,573 total cells (**Figure 1h**, **see Methods**). Although we selected more FOVs from CF samples (CF: 325 FOVs/sample, control: 159 FOVs/sample) and focused on tissue regions in CF individuals with relative cellular preservation to account for the large-scale tissue remodeling, there were still many fewer cells profiled in CF compared to control pancreas (1,646,107 control, 278,466 CF) (**Supplementary Figure 3**). We annotated the cell type identity of spatial gene expression profiles map using scANVI^45^ by first training a reference model on the snRNA-seq data from the single cell multiome map and then applying this model to the spatial map to predict cell type labels (**see Methods**). We further confirmed cell type annotations based on the expression of known marker genes present in the spatial panel (**Figure 1i**). The spatial arrangement of cell types in control individuals matched expectation, while CF showed widescale increases in immune and stellate cell types and very few conventional ductal and acinar cells, mirroring observations in single cell multiome data (**Figure 1j, Supplementary Figure 2c-d**). We also quantified the number of cells per FOV and found that most cell types had decreased numbers per area in CF compared to control, although these decreases were only significant in exocrine cell types (**Supplementary Figure 2e**).

We identified spatial niches and conserved spatial domains across all CF and control samples by clustering and annotating cells based on their neighborhood gene expression profiles using BANKSY^46^ (**Figure 1j-l**). Most niches were primarily composed of exocrine cells, as expected given that >90% of healthy pancreatic tissue is exocrine (**Figure 1k**). Exocrine niches varied widely in their cell type composition, including a niche enriched for stellate, immune, endothelial cells (‘Exo-Stellate’) and a niche enriched for high-enzyme acinar and high-mucin ductal cells (‘Exo-Enzyme’) (**Figure 1k-l**). Niches represented other pancreatic structures as well including islets (‘Endocrine’), comprised primarily of endocrine cells, and large pancreatic ducts (‘Large Ducts’), marked by high-mucin ductal cells (**Figure 1k**). There were marked changes in exocrine-related and other spatial niches in CF including decreased Exo-Acinar, Exo-Ductal and Exo-Stress niches, and increased Exo-Stellate, Exo-Mixed and Immune niches (**Figure 1i**). Overall, this supports the observed large-scale changes to pancreatic composition and structure that occur in CF and highlight specific cell types and spatial structures that are altered.

One of the CF individuals profiled in our study (SCH_CF2) was 2 weeks old, and the spatial organization and proportions of cell types in this donor were extremely different compared to all other CF pancreas samples (which had age range of 14-33) (**Supplementary Figure 4**). Notably, this donor had large amounts of immune cells throughout their pancreas, including an entire tissue region comprised almost entirely of immune cells. It has been well documented that pancreatic destruction and remodeling in CF starts before birth in humans and model organisms^47^ and our observations suggest that widespread inflammation may mark early pancreatic pathogenesis in CF. However, because only a single CF and matched control donor at this age range was profiled, we cannot draw broader conclusions from these observations. Furthermore, as this donor had substantial differences from other CF donors in our study we excluded it from subsequent analyses.

To provide the single cell and spatial profiles generated in this study as a resource to the CF community, we developed an interactive browser which is available at http://cfrdgenomics.org/.

### Pancreatic ductal cells in cystic fibrosis are enriched for a high mucus-producing state

Small pancreatic ducts are among the first structures in the pancreas to break down during CF, as the loss of *CFTR* leads to impaired fluid secretion and ductal clogging^47^. We identified ductal cells in the pancreas of every CF donor, although primarily from the high-mucin ductal population which has lower expression of *CFTR* and normally localize to large pancreatic ducts (**Figure 1g**). High-mucin ductal cells make up a very small proportion of ductal cells in the normal pancreas but showed dramatic increases in CF relative to the conventional ductal population. We confirmed that these cells produce MUC5B protein via immunofluorescence on adjacent tissue sections from the same CF donors (**Figure 2a, Supplementary Figure 5a**). Increased amounts of secreted mucins have been found in CF airway secretions^48^ and likely have a role in CF tissue remodeling; thus, we were interested in characterizing the properties of high-mucin ductal cells further.

**Figure 2.**
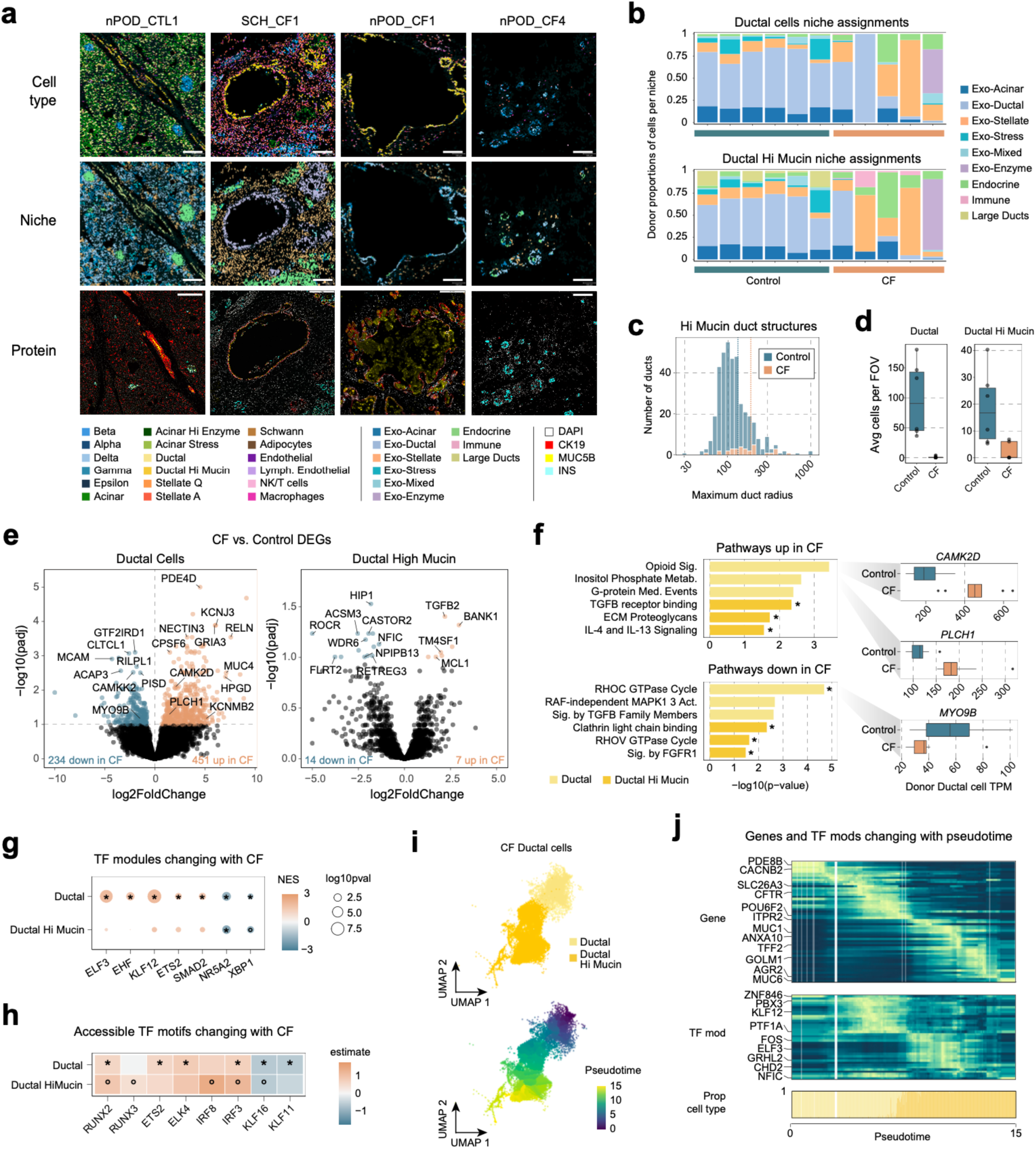
Pancreatic ductal cells transition towards a high mucus-secretion state during cystic fibrosis. a) Example images of four donors profiled with CosMx. Top row: CosMx segmented cells colored by scANVI predicted cell type; middle row: CosMx segmented cells colored by BANKSY niche; bottom row: immunofluorescence images. All immunofluorescence images were from tissue sections from the same donors (consecutive section to the one used for CosMx) and the same regions were used if possible. Scale bars represent 250µm. b) Proportion of CosMx high-mucin ductal cells for each donor classified by CF status (blue=Control, orange=CF) and colored by niche. c) Distribution of radii (µm) of large ductal structures identified using high-mucin ductal cell CosMx spatial coordinates from control (blue) and CF (orange) donors. d) Per donor average number of cells found in an FOV region for both high-mucin and normal ductal cells from Control and CF donors. e) Volcano plots of genes with differential expression in CF donors for both ductal cells (left) and high-mucin ductal cells (right). All genes with differential expression passing FDR<0.1 are colored and the vertical line demarks this cutoff. Genes with increased expression in CF are colored orange and genes with decreased expression in CF are colored blue. f) Left: pathways enriched in genes with significantly increased (top) or decreased (bottom) expression during cystic fibrosis. Bars are colored by the cell type. All pathways have p<0.01 and FDR<0.1 significant enrichments are marked with *.Right: distribution of per-donor ductal cell TPM values for select significantly up- or down-regulated genes from enriched pathway leading edge gene lists. g) TF modules with target genes that are significantly upregulated or downregulated in cystic fibrosis. The color of each dot denotes the normalized enrichment score (NES) value from fGSEA and the dot size represents the -log10 of the p-value for the pathway enrichment. FDR<0.1 significant enrichments are marked with * and p<0.01 are marked with o. h) Heatmap of motifs with increased or decreased global accessibility in cell types during CF. Cells are colored by a linear mixed model estimate for the effect of cystic fibrosis. FDR<0.1 significant enrichments are marked with * and p<0.01 are marked with o. i) UMAP of all ductal cells from CF donors colored by cell type (top) or pseudotime (bottom). j) Genes and TF modules with per-cell expression or enrichment correlated with pseudotime.

In the non-diseased pancreas, high-mucin cells were primarily found in large pancreatic duct structures (Large Ducts niche) although they were also present in other exocrine niches (**Figure 2a-b**). High-mucin ductal cells across different niches had distinct gene expression profiles, likely reflecting different functions and cellular environments. For example, high-mucin ductal cells within large pancreatic ducts were significantly enriched (FDR<.10) for genes involved in cell-cell adhesion (e.g. ephrin receptor binding, cadherin binding), interactions with immune cells (e.g. cytokine receptor binding, chemokine activity) and protein processing (endopeptidase regulator activity) (**Supplementary Figure 5b**). Conversely, high-mucin ductal cells in the Exo-Ductal niche had higher expression of genes involved in the response to growth factors and interactions with proteases, likely due to neighboring acinar cells in this niche (**Supplementary Figure 5b**).

In the CF pancreas, high-mucin ductal cells had altered localization and niche composition compared to control pancreas. High-mucin cells were in both large and small duct structures in partially remodeled tissue in CF and rarely present in highly remodeled regions (**Figure 2a**), and high-mucin cells in these structures mapped to different niches. Additionally, high-mucin cells were less abundant in the main exocrine (Exo-Acinar, Exo-Ductal) niches and, conversely, were substantially more abundant in the Exo-Stellate and Exo-Enzyme niches (**Figure 2b**). We clustered high-mucin ductal cell spatial coordinates to identify distinct duct structures and found a large decrease in the number of ducts in CF (**Figure 2c**), reflecting a broad breakdown of ductal structures. Additionally, while the number of high-mucin ductal cells per spatial region (FOV) decreased in CF, this reduction was much less pronounced compared to the conventional ductal population (**Figure 2d**). This demonstrates that high-mucin cells are relatively preserved in CF compared to conventional ductal cells yet with altered niche composition, which may reflect greater ability of these ductal cells to adapt to different tissue environments.

We next characterized CF-associated changes in gene expression and regulation in both the conventional ductal and high-mucin ductal populations. Although ductal and high-mucin ductal cells had comparable numbers of cells in the single cell map (19,045 ductal, 15,076 high-mucin ductal), there were dramatically more changes within the conventional ductal population in CF (**Figure 2e-h**). Hundreds of genes had significantly increased expression (FDR<.10) in ductal cells in CF, including genes involved in ion sensing and transport (*KCNJ3*, *CAMK2D*, *KCNMB2*) and *PDE4D*, a gene previously implicated in *CFTR* regulation^49^ (**Figure 2e, Supplementary Table 5**). Cellular processes including G-protein mediated events and inositol phosphate metabolism were upregulated in ductal cells in CF (**Figure 2f, Supplementary Table 6**), which form a broader pathway leading to the release of Ca^2+^ ions^50^ that can facilitate HCO_3-_ secretion in the absence of *CFTR*^51^. Ductal cells also downregulated genes involved in Rho GTPase signaling and TGFβ signaling (**Figure 2f**), where Rho GTPases are involved in the actin cytoskeleton and vesicle trafficking^52^ and TGFβ in response to stress, proliferation and extracellular matrix remodeling^53^. These results illustrate adaptation of ductal cells to loss of *CFTR* function in CF, including impaired cytoskeleton trafficking, reduced proliferation, and higher stress responses that may accompany ductal cell death in CF.

High-mucin ductal cells were marked by upregulation of secreted mucins and relatively low expression of *CFTR* compared to the conventional ductal population^54^ (**Supplementary Figure 5c**). These cells are thus likely less influenced directly by mutations in *CFTR* and accordingly show less dramatic changes in CF compared to conventional ductal cells. Despite this, high-mucin ductal cells in CF did show evidence for changes in several cellular processes including those involved in cell structure, stress, and secretion. For example, CF high-mucin ductal cells had increased expression (FDR<.10) of pathways involved in ECM remodeling (ECM proteoglycans, molecules associated with elastic fibers) and TGFβ signaling, and decreased expression of pathways involved in mitochondrial metabolism and vesicle trafficking (**Figure 2f**). Overall, these results suggest that the high-mucin ductal cells undergo metabolic changes that may promote survival and potentially even contribute to large-scale tissue fibrosis in CF.

To identify regulatory networks driving changes in ductal cell expression in CF, we identified TF GRN modules with altered expression in ductal cells in CF (**Supplementary Table 7**) as well as TFs with global changes in sequence motif accessibility in CF (**Supplementary Table 8**). As with gene expression changes, there were overall more changes in TF activity in the conventional ductal population compared to high-mucin ductal cells (**Figure 2g-h**). There was increased activity of ETS family TFs (*ELF3*, *EHF*, *ETS2*, *ELK4*) in conventional ductal and high-mucin ductal cells, which likely facilitate the increased secretion and stress responses found in these cells. In addition, we identified increased activity of TFs such as IRF and RUNX which regulate stress and fibrosis (**Figure 2h**). Additionally, the GRN for SMAD2, a TF directly downstream of TGFβ signaling, was significantly upregulated in ductal cells (**Figure 2g**). Finally, we found a decrease in the expression of GRNs for TFs that regulate pancreatic identity and metabolism, and *XBP1*, a master regulator of ER stress (**Figure 2g**), which may help drive the decreased activity of metabolic and secretion pathways in ductal cells.

Interestingly, we found evidence for changes in the activity of multiple *KLF* family TFs, which are involved in pancreatic exocrine cell differentiation and proliferation^55^. There was increased activity of *KLF12* in ductal cells in CF and decreased activity of other KLF family TFs including *KLF16* and *KLF11* (**Figure 2g,h**). *KLF11* mediates the antiproliferative effect of TGFβ on pancreatic cell types^56^ whereas *KLF12* is a transcriptional repressor which can antagonize and prevent the activity of other KLF factors^57^. While *KLF12* showed expression in both ductal and high-mucin ductal cells in the CF pancreas, *KLF11* specifically had much lower expression compared to control pancreas (**Supplementary Figure 5d**). Together these indicate an overall decrease in the regulation of genes by KLF family TF in ductal cells in CF which may help ductal cells survive under stressful conditions in CF. Among cREs with significantly altered accessibility (FDR<0.1) in CF ductal cells (**Supplementary Table 9**), we highlighted a cRE containing a *KLF12* motif and predicted to regulate the expression of *NCALD* (**Supplementary Figure 5e**). The *NCALD* gene encodes a calcium sensor protein involved in endocytosis, which supports that *KLF12* activity may drive changes in ion sensing and signaling in ductal cells during CF.

Marker genes of high-mucin ductal cells were significantly upregulated in the conventional ductal population in CF (**Supplementary Figure 5f**), suggesting a potential transition to the high-mucin subtype. To investigate a state transition of ductal cells in CF, we performed trajectory analysis starting from the conventional ductal population and calculated pseudotime for all ductal cells in CF donors (**Figure 2j**). We then identified genes and TF modules correlated with progression along the pseudotime axis across cells (**Figure 2k, Supplementary Table 10**). As expected, the trajectory was anchored by the expression and activity of markers of conventional ductal identify (*CFTR, PBX3*) and secretory processes (*PDE8B, CACNB2*) on one end, and markers involved in mucus secretion (*MUC6*, *AGR2*) and chromatin remodeling (*CHD2, NFIC*) on the other. Genes in the middle of the trajectory that may represent points of transition include the pancreatic developmental TF *POU6F2* and genes and TFs involved in cell stress (*ANXA10*, *TFF2*, *GOLM1*, *FOS, ELF3*). Additionally, *ITPR2*, an IP3 receptor involved in Ca^2+^ ion release, and the *KLF12* GRN were correlated with pseudotime and peaked right before the transition to high-mucin ductal cells (**Figure 2k**), This provides further evidence for specific factors that may facilitate changes in ductal cell ion channel activity and secretion during CF which may lead ductal cells towards a mucus secretion-related phenotype.

### Inflammatory and pro-fibrotic cell communication facilitates increased activity in CF

In addition to changes in ductal cell sub type compositions, we also identified large increases in the proportions of inflammatory, stellate, and endothelial cell types in the pancreas of CF donors (**Figure 1g**). In the healthy pancreas, immune cells including macrophages, T and NK cells were found around the large ducts and scattered throughout the exocrine tissue in regions marked by the Exo-Stellate niche, which had increased abundance in CF donors (**Figure 1g**, **Figure 3a**). Pro-fibrotic activated stellate cells and quiescent stellate cells were found in similar locations and the same niches with endothelial and lymphatic endothelial cells (**Figure 3a**). In the CF pancreas, however, both activated and quiescent stellate cells were found throughout the tissue and were often very close to islets within large clusters of immune cells (**Figure 3a**).

**Figure 3.**
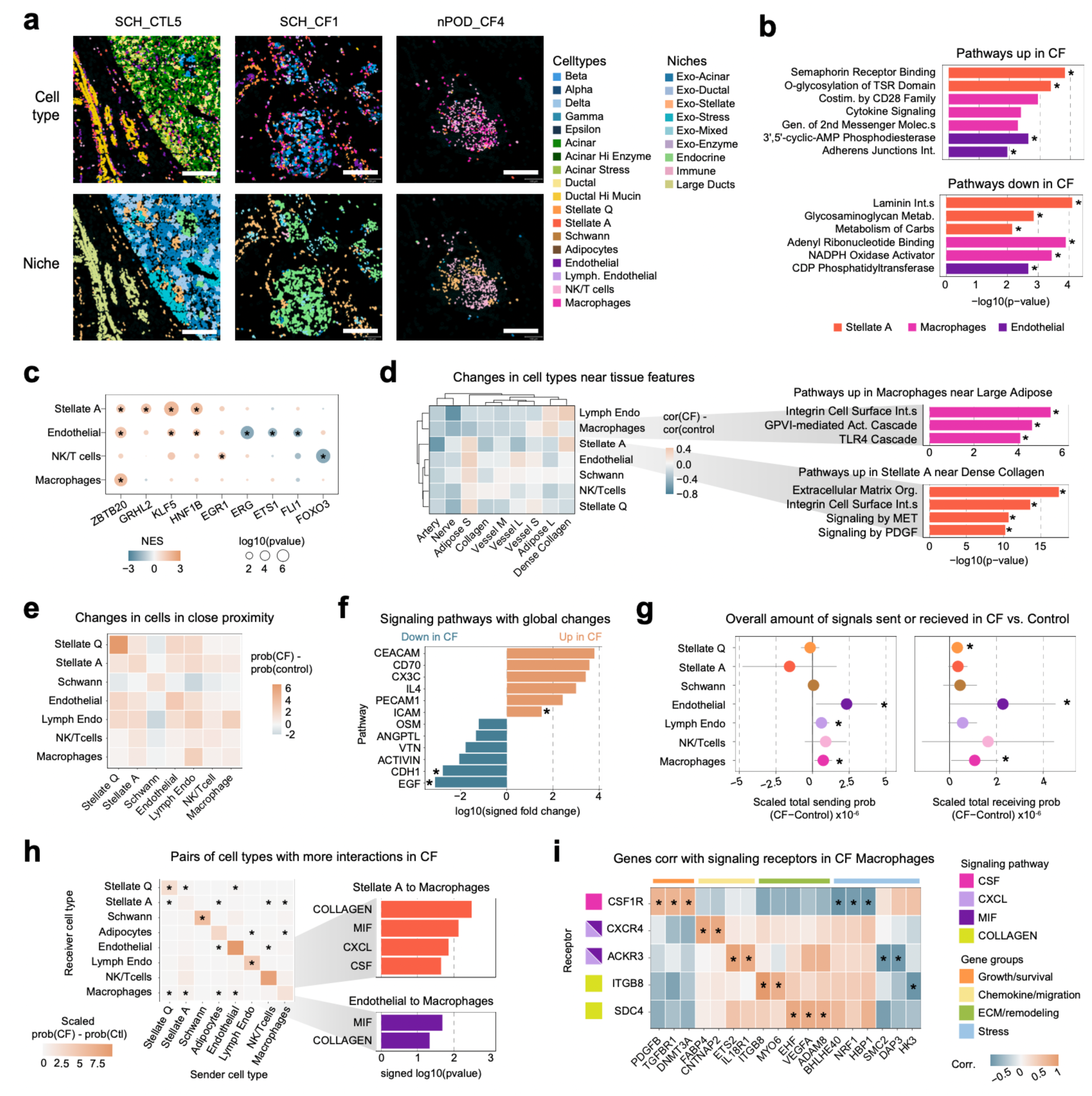
Crosstalk between pro-fibrotic and inflammatory cell types drives increased activity in CF. a) Example images of three pancreata profiled with CosMx. Top row: CosMx segmented cells colored by scANVI predicted cell type; bottom row: CosMx segmented cells colored by BANKSY niche. Scale bars represent 200µm. b) Pathways enriched in genes with significantly increased (top) or decreased (down) expression during cystic fibrosis. All pathways have p<0.01 and FDR<0.1 significant enrichments are marked with *. c) TF modules with target genes that are significantly upregulated or downregulated during cystic fibrosis. The color of each dot denotes the normalized enrichment score (NES) value from fGSEA and the dot size represents the -log10 of the p-value. FDR<0.1 significant enrichments are marked with *. d) Left: changes in cell type proximity to tissue feature annotations in CF. Correlation tests were run using the proportion of a cell type at increasing distances from a tissue feature and then multiplied by -1. The plot represents the difference in correlation between CF and control cells. Right: Pathways enriched in genes that had increased expression with proximity to a tissue feature in CF cells. FDR<0.1 significant enrichments are marked with *. e) Difference in the probability of pairs of cell types being in close proximity (<50µm) between CF and control pancreata. f) Cell-cell signaling pathways with increased or decreased signaling probability between any cell types in CF or control pancreata. Fold change values are signed based on the direction of change, where the FC of pathways increased in CF are positive and the FC of pathways increased in control are negative. Pathways with nominally significant (p<0.05) changes in per-donor average signaling levels are marked with *. g) Differences in the scaled total probability of signals sent (left) or received (right) by a cell type in CF vs. Control. Points represent the difference between CF and control per-donor averages across all pathways, 95% confidence intervals are plotted as bars and p<0.05 is marked with *. h) Left: pairs of cell types with more signals sent in CF than control. The average signaling probability between each pair of cell types was calculated per donor and then a t-test was run to identify differences between CF and control. Cells are colored by the cell type size and z-score scaled difference in probability, as calculated on all pairs of cell types. Pairs of cell types with p<0.05 increased interactions are marked with *. Right: pathways with p<0.05 increased signaling probability between a pair of cell types in CF donors. i) Rank-based correlation of per-donor expression of signaling receptors with genes in CF macrophages. FDR<0.1 correlations are marked with *.

Gene expression and chromatin profiles from immune, stellate, and endothelial cells supported increased activity in disease. Within these cell populations, activated stellate cells and macrophages had the most significant changes in gene expression in CF (**Supplementary Figure 6a, Supplementary Table 5**). At the pathway level, activated stellate cells had significantly increased expression (FDR<.10) of semaphorin receptor binding and TSR protein glycosylation, and decreased laminin interactions and glycosaminoglycan metabolism (**Figure 3a, Supplementary Table 6**). Together this indicates that activated stellate cells are shifting towards a migratory phenotype that is actively remodeling matrix and promoting fibrotic collagen formation over laminin, a component of the basement membrane in the healthy pancreas^58^. Target genes of TFs known to promote fibrosis in the pancreas, including *ZBTB20*^59^ and *KLF5*^60^, had significantly increased expression (FDR<.10) in activated stellate cells in CF (**Figure 3c, Supplementary Table 7**). Macrophages also had increased activity of target genes of pro-inflammatory *ZBTB20* (**Figure 3c, Supplementary Figure 6b**) and increased expression of immune activation and inflammatory pathways (e.g. cytokine signaling, co-stimulation by CD28) (**Figure 3b**). NK and T cells had significantly increased expression (FDR<.10) of target genes for EGR1, a factor involved in early activation of T cells that drives cytokine production^61^ (**Supplementary Figure 6c**) and decreased expression of target genes for FOXO3, which promotes NK cell development^62^ (**Figure 3c**).

The increased expression of activation-related pathways in immune and fibrotic cell types was especially pronounced when considering the spatial context of these cells. For example, activated stellate cells were generally proximal to regions of dense collagen in the CF pancreas compared to controls (**Figure 3d, Supplementary Table 11**). Cells closer to regions of dense collagen had significantly increased expression (FDR<.10) of genes involved in matrix formation and interactions (e.g. extracellular matrix organization, integrin cell surface interactions, signaling by PDGF), as well as pathways that promote cell proliferation (signaling by MET) (**Figure 3d, Supplementary Table 12**). Conversely, macrophages were more proximal to large adipose tissue in CF and cells closer to these regions had increased activity (FDR<.10) of pathways involved in production of pro-inflammatory cytokines and matrix interactions (**Figure 3d**).

Compared to inflammatory and fibrotic cell types, endothelial cells had decreased expression (FDR<.10) of target genes of multiple transcription factors that maintain endothelial cell identity^63^ (*ERG*, *ETS1*, *FLI1*), as well as genes involved in maintenance of cell membrane integrity (**Figure 3b-c, Supplementary Figure 6d**). Endothelial cells in CF had increased expression (FDR<.10) of genes involved in the breakdown of cyclic-AMP, an intracellular messenger that facilitates anti-inflammatory processes and cell-cell interactions (e.g. adherens junction interactions) (**Figure 3b**). Additionally, the target genes of *KLF5*, associated with vascular remodeling and endothelial dsyfunction^64^, were upregulated in CF (**Figure 3c**). Overall, these findings are consistent with dysfunction and remodeling of endothelial cells in the pancreas in CF.

To interrogate cellular interactions driving changes in immune, stellate and endothelial cells in CF, we examined changes in cell-cell proximity and signaling in CF compared to control pancreas (**Supplementary Data 2**). Almost all inflammatory, stellate, and endothelial cells showed closer proximity (<50µm) to themselves and each other in CF (**Figure 3e, Supplementary Table 13**). At a global level, multiple cell-cell signaling pathways had nominally significant changes in interactions in CF (p<0.05). Specifically, the intracellular adhesion molecules (ICAM) pathway had increased signaling in the CF pancreas, while the epidermal growth factor (EGF) and cadherin (CDH) pathways had decreased signaling (**Figure 3f, Supplementary Table 14**). These reflect an overall increase in fibrotic signaling in CF and a decrease in normal exocrine driven signaling. Among non-exocrine and non-endocrine cell types, macrophages, and endothelial cells both received and sent more signals in CF, lymphatic endothelial cells sent more signals, and quiescent stellate cells received more signals (**Figure 3g, Supplementary Figure 6e, Supplementary Table 15**).

Many pairs of inflammatory, fibrotic, and endothelial cells had increased probability of signaling in CF compared to control (**Figure 3h, Supplementary Table 16**). For example, macrophages received increased signals from activated stellate and endothelial cells, including increased collagen and many signals known to recruit and sustain macrophage activity (MIF, CXCL, CSF) (**Figure 3h, Supplementary Table 17**). We leveraged single cell multiome profiles to predict the downstream effects of signaling within macrophages by correlating signaling receptor expression with gene and TF module activity (**see Methods**). The expression of *CSF1R*, the primary receptor for macrophage colony-stimulating factor (CSF), was correlated with the expression of genes promoting cell growth and survival in CF individuals (**Figure 3i, Supplementary Table 18**). Receptors for CXCL and MIF signals were correlated with genes promoting chemokine release and migration, and receivers of collagen signaling via *ITGB8* and *SDC4* were correlated with genes involved in ECM production and remodeling (**Figure 3i, Supplementary Figure 6f**). By comparison, all of these signaling receptors in macrophages were inversely correlated with genes involved in stress and anti-inflammatory pathways, further supporting that signals are generally associated with increased macrophage activity (**Figure 3i, Supplementary Figure 6f**).

### Endocrine cells in CF show marked changes distinct from other forms of diabetes

The majority of endocrine cells persist in the pancreas in CF despite large scale tissue remodeling, although impaired glucose tolerance is common in individuals with CF and most develop CFRD^32^. We identified many genes in both beta and alpha cells with altered gene expression profiles in CF compared to control (**Figure 4a, Supplementary Table 5**). Surprisingly, in beta cells, many genes upregulated in CF were involved in secretion-related processes, including *RAB11A*, *HCN1*, and *EXOC1*, and multiple pathways related to insulin secretion also had increased activity in CF (**Figure 4a-b, Supplementary Table 6**). In line with this, TFs linked to beta cell identity and insulin secretion such as *ISL*, *PAX6* and *NEUROD1* had increased accessibility (FDR<.10) in CF beta cells and increased expression of TF target genes (FDR<.10) (**Figure 4c-d, Supplementary Tables 7-8**). Overall, these changes indicate that beta cells in CF may show increased insulin secretion, potentially as a compensatory mechanism. Alpha cells in CF had increased activity of multiple calcium channel genes (*CACNA2D1, CACNA2D3*), suggesting the potential for increased secretory activity as well (**Figure 4b**).

**Figure 4.**
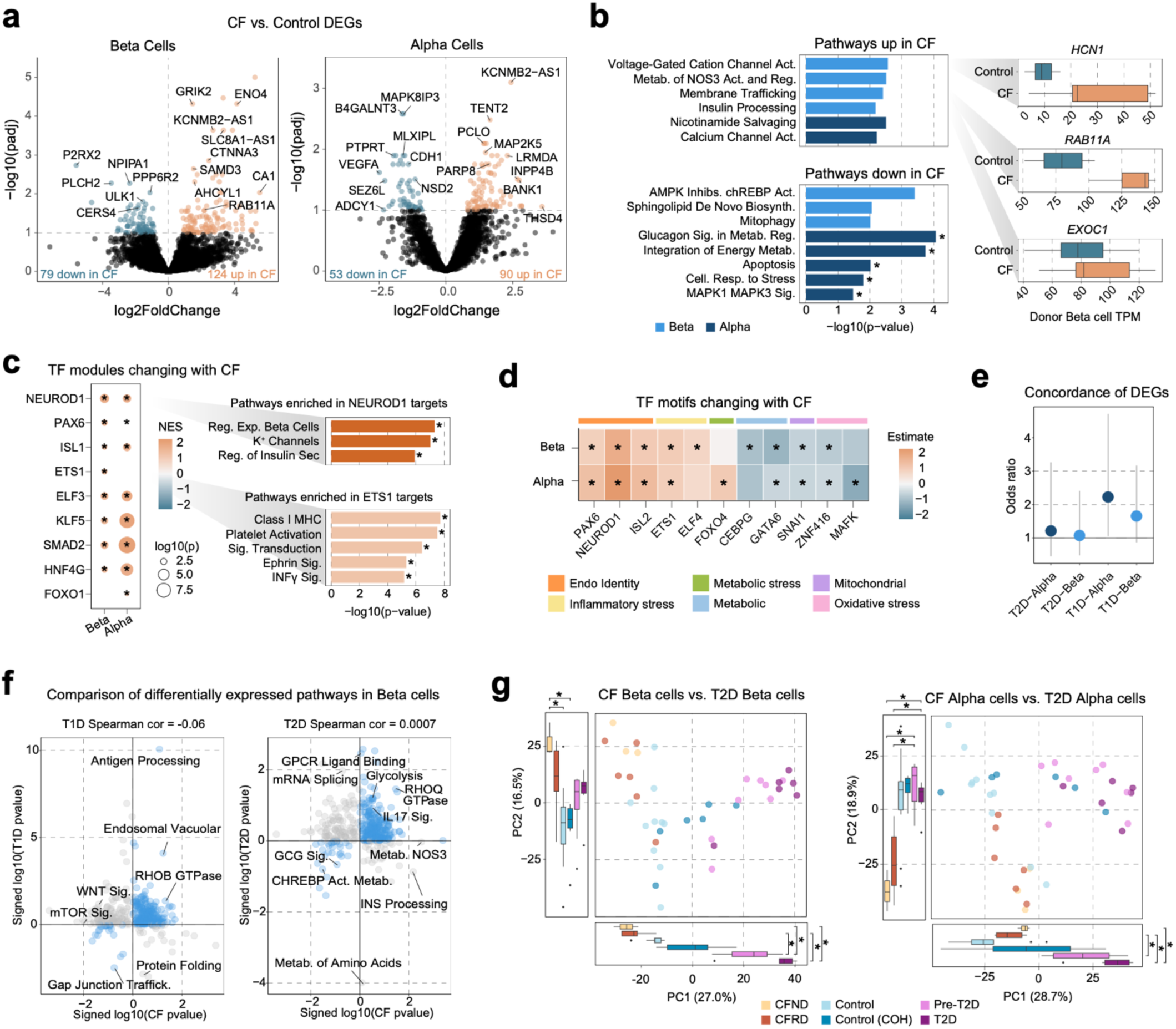
Changes in beta and alpha cell genomic profiles in CF are distinct from type 1 and type 2 diabetes. a) Volcano plots of genes with differential expression in CF donors for both beta cells (left) and alpha cells (right). All genes with differential expression passing FDR<0.1 are colored and the horizontal line demarks this cutoff. Genes with increased expression in CF are colored orange and genes with decreased expression in CF are colored blue. b) Left: pathways enriched in genes with significantly increased (top) or decreased (bottom) expression during cystic fibrosis. All pathways have p<0.01 and FDR<0.1 significant enrichments are marked with *. Right: distribution of per-donor TPM values for select significantly up-regulated genes from beta cells. c) Left: TF modules with target genes that are significantly upregulated during cystic fibrosis. The color of each dot denotes the normalized enrichment score (NES) value from fGSEA and the dot size represents the -log10 of the p-value. FDR<0.1 significant enrichments are marked with *. Right: pathways enriched in the target genes of select TFs. d) Heatmap of motifs with increased or decreased global accessibility in cell types during CF. Cells are colored by a linear mixed model estimate for the effect of cystic fibrosis. The colored bars above the heatmap indicate broad annotations of the cellular processes a TF is involved in. FDR<0.1 significant enrichments are marked with *. e) Gene set concordance between genes upregulated with CF in beta and alpha cells and genes upregulated with T1D or T2D in the cell type. f) Comparison of pathways up- and down- regulated in CF beta cells with changing pathways in T1D and T2D beta cells. P-values are signed based on the direction of effect and pathways passing FDR significance in either or both studies are colored. g) PCA comparison of donor pseudobulk RNA profiles with pre-T2D and T2D donors from an external dataset. A type 2 ANOVA and then a Tukey’s test were run to identify groups of donors with different mean PC embeddings. All p<0.05 differences between a CF group and a T2D group are annotated with *.

Beta cells in CF also had increased expression of genes involved in stress signaling (**Figure 4b**), and both beta and alpha cells had increased expression of target genes for stress-response related TFs including inflammatory stress (*ETS1*, *ELF3*, *ELF3*) and metabolic stress (*KLF5*, *HNF4G*, *SMAD2*) (**Figure 4c-d**). Target genes of *ETS1* were enriched (FDR<.10) in pathways involved in inflammatory signaling and cellular interactions (antigen presentation folding, assembly and peptide loading of class I MHC, interferon gamma signaling) and signaling pathways (signal transduction, Eph-Ephrin signaling), suggesting the elevated stress responses are driven in part by external factors (**Figure 4c**). Alpha cells specifically had significantly increased (FDR<.10) expression of targets of FOXO-family TFs and increased accessibility of FOXO motifs in CF (**Figure 4c-d**). The FOXO family of TFs are responsible for facilitating cellular responses to stress and have been found to increase glucagon secretion^65^, indicating that alpha cells may have increased function in response to stress in CF.

The increase in cellular stress in CF is also reflected in the overall downregulation of pathways involved in metabolic functions in both cell types (e.g. beta cells - AMPK Inhibits chREBP transcriptional activation activity, sphingolipid de novo biosynthesis, mitophagy; alpha cells - glucagon signaling in metabolic regulation, integration of energy metabolism) (**Figure 4b**). Beta and alpha cells both also had significantly decreased accessibility of multiple families of TF motifs known to promote metabolic processes (beta cells: *CEBPG*; alpha cells: *GATA6*) (**Figure 4c-d**). Finally, there was significantly reduced (FDR<.10) expression of proliferation-related pathways in CF alpha cells (MAPK1/MAPK3 signaling). Taken together, these results reveal increased stress responses and reduction in metabolic processes in beta and alpha cells in CF.

The pathogenesis of CFRD has similarities to both type 1 diabetes (T1D) and type 2 diabetes (T2D), and genetic risk factors for T2D also overlap CFRD risk^66^ (**Supplementary Figure 7a**), but the relationship in genomic changes between CFRD, T1D and T2D is largely unknown. We compared genes with altered activity in CF to those with altered activity in T1D and T2D from a previous scRNA-seq study in islets^44^. There was minimal overlap in genes altered in CFRD and T1D/T2D in both cell types with slightly higher concordance with T1D, and no estimate was significant (**Figure 4e, Supplementary Table 19**). We further compared cellular pathways with enriched expression in CF, T1D and T2D and there was a similar lack of overlap (**Figure 4f**). Several individual processes had similarities between CF and T1D or T2D in beta cells, including increased antigen processing in both CF and T1D and decreased GCG signaling in both CF and T2D. However, many cellular processes altered in beta cells in T1D or T2D did not have significant, directionally consistent changes in CF beta cells, and vice versa (**Figure 4f**). Furthermore, PCA of gene expression profiles from CF, T1D and T2D donors highlighted largely distinct profiles between forms of diabetes (**Figure 4g, Supplementary Figure 7b**).

### Proximity to adipose tissue and collagen drives beta cell dysfunction and loss in CF

To understand changes in individual islets in CF, we defined islets in each FOV using the concave hull created by the spatial coordinates of endocrine cells (**see Methods**) (**Figure 5a, Supplementary Data 3**). Islets in CF had significant structural and compositional differences compared to control (**Figure 5b, Supplementary Table 20**). On average, islets in CF had fewer cells and smaller area and were denser than control islets. Islets in CF also had a higher proportion of bordering (peri-islet) activated stellate cells, macrophages and endothelial cells. Further, islets in CF were more likely to be located near large regions of adipose tissue (>100,000µm^2^), which are the result of large-scale adipose accumulation that occurs in CF pancreas due to tissue remodeling. Endocrine cell types had increased probability of being proximal (<50µm) to regions of large adipose tissue in CF compared to control (**Figure 5c, Supplementary Table 11**). Similarly, endocrine cell types had increased probability of being in proximity (<50µm) to collagen deposits, another hallmark feature of the pancreas in CF.

**Figure 5.**
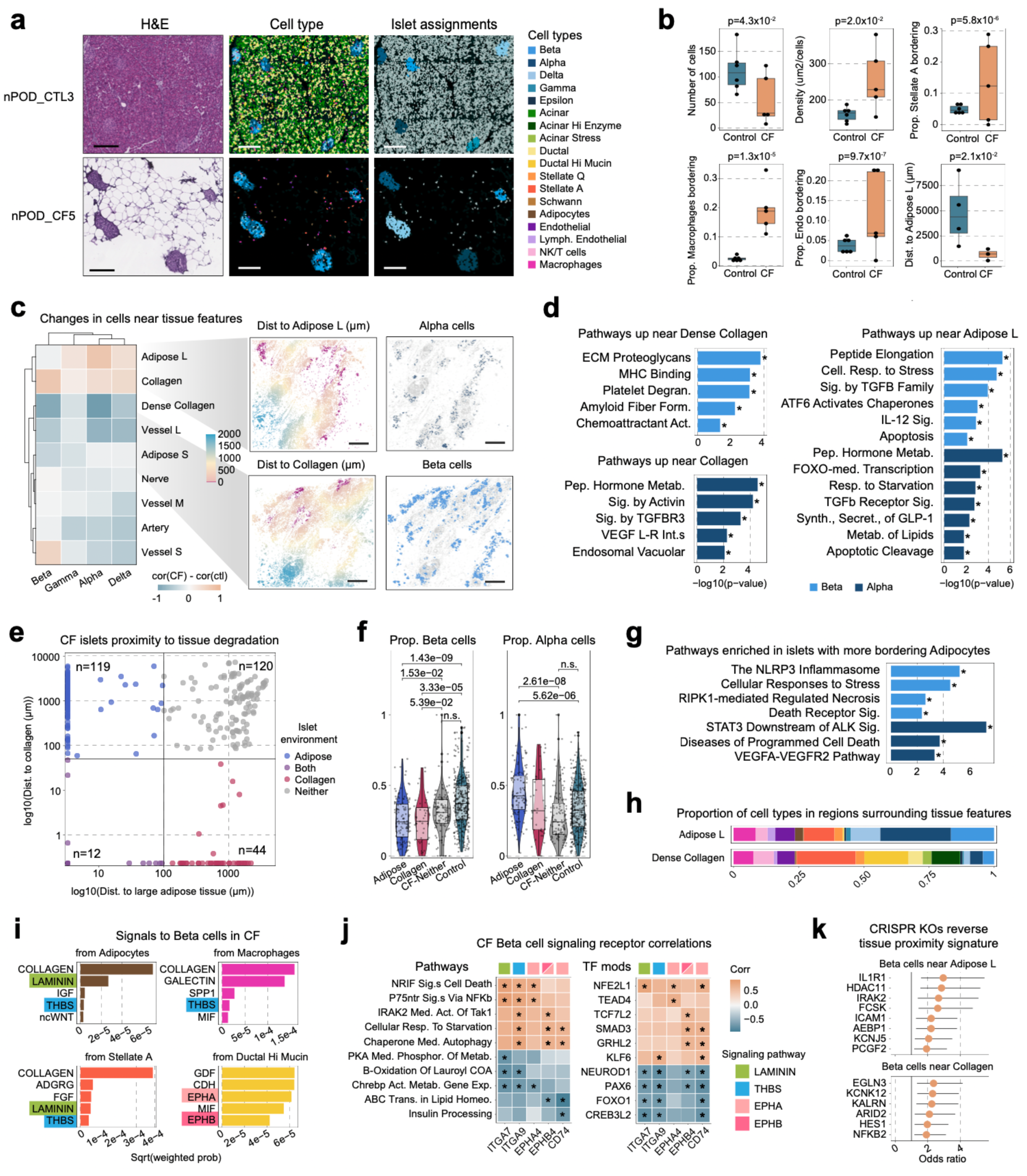
Proximity to large adipose tissue in CF pancreas drives beta cell stress and death. a) Example images of regions with islets from two pancreata profiled with CosMx. Left: H&E stain; middle: CosMx segmented cells colored by scANVI predicted cell type; right: CosMx segmented cells colored by specific islet assignments (blues) or no assignment (grey). Scale bars represent 300µm. b) Donor averaged per-islet measurements with significant (t-test p<0.05) differences between control and CF. c) Left: changes in cell type proximity to tissue feature annotations in CF. Correlation tests were run using the proportion of a cell type at increasing distances from a tissue feature and then multiplied by -1. The plot represents the difference in correlation between CF and control cells. Right: example regions of CF pancreata colored by proximity to a tissue feature (left) and specific cell type (right). Scale bars represent 1,000µm. d) Pathways enriched in genes that had increased expression with proximity to a tissue feature in CF cells. FDR<0.1 significant enrichments are marked with *. e) Distribution of distance to large adipose tissue and collagen annotations among all islets from annotated CF donors. Environment was classified based on whether an islet was within 100µm of large adipose tissue and/or 50µm of collagen. f) Per-islet proportion of beta (left) and alpha (right) cells. Significant comparisons from a type 2 ANOVA and Tukey comparison are marked with bars and annotated with the p-value. g) Pathways enriched in genes that had increased expression in islets with higher proportion of bordering Adipocytes. FDR<0.1 significant enrichments are marked with *. h) Proportions of cell types in all cells surrounding large adipose tissue (<100µm) and dense collagen (<50µm). i) The top 5 signals to beta cells in CF from select cell types. The average p-value weighted signaling probability across all ligand-receptor interactions per pathway were plotted. Only signaling pathways where the sender cell type had an average TPM>10 for pathway ligands are included. j) Rank-based correlation of per-donor expression of signaling receptors with average pathway enrichments in CF beta cells. FDR<0.1 correlations are marked with *. k) Gene set concordance between genes upregulated in CF beta cells near large adipose tissue or collagen and genes with decreased expression after CRISPR knockout of a select gene (y-axis).

Both beta and alpha cells had higher expression of genes involved in stress response pathways with increased proximity to collagen and adipose features in CF (**Figure 5d, Supplementary Table 12**). For example, near dense collagen, beta cells significantly up-regulated (FDR<.10) the expression of genes involved in fibrosis response, amyloid fiber formation and immune interactions (**Figure 5d**). Alpha cells near collagen in CF had increased expression of genes involved in signaling responses, as well as secretion and vascularization (**Figure 5d**). Alpha cells near large adipose up-regulated stress pathways as well as processes involved in promoting function and survival during stress, indicating that alpha cells may be adapting to stress by increasing hormone secretion. For example, FOXO-mediated transcription was increased in alpha cells near adipose tissue, as well as multiple hormone synthesis and secretion pathways (**Figure 5d**). Conversely, beta cells near large adipose tissue upregulated multiple stress related pathways as well as cell death processes (**Figure 5d**). This suggests that beta cells may show less ability to adapt to stress in specific tissue environments such as large adipose regions in CF.

We classified individual islets from CF donors based on their proximity to both large adipose and collagen regions (**Figure 5e**). Islets from CF donors were primarily located either near adipose tissue (<100µm, n=119), collagen (<50µm, n=44) or neither (n=120), with few islets found near both features (n=12) (**Figure 5e**). CF islets in large adipose tissue had a significantly lower proportion of beta cells and higher proportion of alpha cells, compared to islets not in direct proximity to either feature or islets from control pancreas (**Figure 5f, Supplementary Table 21**). Islets in collagen also had significantly reduced proportions of beta cells, but with no change in the proportion of alpha cells. By comparison, islets not proximal to either large adipose or collagen had no significant change in beta cell proportion compared to control pancreas islets. Beta cells in islets with a higher proportion of surrounding adipocytes based on the spatial annotations had significantly increased (FDR<.10) expression of stress response and cell death pathways (**Figure 5g**). Alpha cells in CF islets with more bordering adipocytes also had increased expression of cell death pathways but also cell survival and vascularization pathways (**Figure 5g**).

We identified specific signaling interactions in large regions of adipose tissue and collagen driving beta cell death in CF. We first examined the cell type composition of these regions, which included adipocytes, macrophages, activated stellate cells, and high-mucin ductal cells (**Figure 5h**). Next, we identified incoming signals to beta cells from these cell types, which included matrix-interaction (collagen, laminin, thrombospondin), inflammatory (galectin, MIF, CXCL) and proliferation (IGF, WNT, TGFβ) signals (**Figure 5i**). We correlated the expression of receptors for each signal to pathway expression in beta cells to predict the downstream effects of these signaling interactions (**see Methods, Supplementary Table 22**). Receptors involved in cell-matrix signaling *ITGA7*, *CD47*, and *EPHA3* were significantly correlated with cell stress and cell death pathways (**Figure 5j**). Additional ephrin receptors *EPHA4* and *EPHA5* were significantly correlated with dysregulated cellular metabolism pathways (**Figure 5j**). Conversely, the expression of *ITGA7*, *CD47*, and *EPHA3* inversely correlated with metabolic pathways, and the expression of ephrin receptors inversely correlated with pathways involved in lipid homeostasis and insulin processing (**Figure 5j**).

Given genomic changes in beta cells proximal to adipose tissue and collagen linked to dysfunction and cell death in CF, we finally prioritized small molecules and gene knockouts that may represent targets to reverse these effects. We identified gene sets up- or down-regulated by small molecule exposure or CRISPR gene knockout from the LINCS database that showed evidence (p<.01) for inverse concordance to genes in beta cells with increased or decreased expression near adipose or collagen structures (**Supplementary Table 19**). For example, based on effect size, genes altered by knockout of the cytokine receptor *IL1R1*, immune response gene *IRAK2,* histone deacetylase *HDAC11,* and cell surface protein *ICAM1* showed strongest inverse overlap with genes with altered expression in beta cells near large adipose tissue (**Figure 5k**). Thus, these molecules and genes may represent candidates to reverse beta cell dysfunction and prevent beta cell loss due in CF.

## Discussion

Single cell and spatial profiling of the pancreas from individuals with CF provided unprecedented insight into changes in the pancreas that drive the development of CFRD. While there is considerable damage to pancreatic ductal structures and loss of ductal cells in CF^12,47,67^, we unexpectedly identified a population of ductal cells marked by high expression of secreted mucins that was relatively preserved in CF. High-mucin ductal cells had been previously identified in the healthy pancreas by single cell and imaging studies, where they localize to large ducts and pancreatic ductal glands^35,43,54^, although with very low abundance overall. The preservation of high-mucin ductal cells in CF may contribute to the overall tissue pathology seen in CF in beyond physical obstruction of the ducts. Studies in other tissues have found that secreted mucins including *MUC5B,* a key marker of the high-mucin cells, can trigger inflammation^68^ which could contribute to pancreatic damage in CF. In the present study, high-mucin ductal cells showed upregulated pro-fibrotic processes and decreased epithelial integrity and cell-cell junctions, along with dramatically altered tissue localization and niche composition in the CF pancreas. Therefore, these cells may promote tissue remodeling and fibrosis throughout the pancreas in CF. Furthermore, high-mucin ductal cells in areas of tissue damage demonstrated altered signaling to beta cells, and thus may contribute directly to the beta cell dysfunction observed in CF. The small number of conventional ductal cells that remained in CF up-regulated processes active in high-mucin cells and thus may transition to the high-mucin sub-type, which could further exacerbate disease. Finally, the altered activity in ductal cells of KLF TFs, which have been implicated in ductal adenocarcinoma^71^, will require additional investigation given the concern of increased relative risk of pancreatic cancer observed in individuals with CF.

Beta cell dysfunction and reduced beta cell mass are hallmarks of all forms of diabetes, but the extent of changes in beta cells in CF are relatively unknown. Our study confirmed morphological differences of islets in CF including smaller islet area and increased immune infiltration^22,23,31^, as well as overall reduced beta cell mass^22,23^. In contrast to previous studies which found minimal change in key beta cell genes in whole-islet transcriptomes in CF^23^, we identified marked genomic changes in beta cells in CF, suggesting beta cell dysregulation is also a feature of CF. We also uncovered novel insight into the drivers of beta cell loss in CF. There was a significantly decreased proportion of beta cells in islets within regions of large adipose and collagen deposition in CF where, by comparison, islets in CF outside these areas showed no difference in beta cell proportion from non-disease. Proximity to areas of tissue damage was also strongly correlated with increased activity of stress and cell death-related pathways in beta cells. Thus, large adipose and collagen deposits likely represent major contributors to beta cell dysfunction and loss in CF. The mechanisms driving this loss may involve signaling interactions from the specific cell types in damaged areas, such as high-mucin ductal, stellate, adipose and macrophage cells, as these signals were linked to downstream increases in stress and cell death pathways in beta cells. Compared to beta cells, alpha cells were relatively preserved in CF, as has been reported previously^22,23,31^. Therefore, despite being localized to the same areas of tissue damage and showing similar increases in stress responses, alpha cells may have greater ability to adapt and survive during stress, as has been described in T1D and T2D.

Notably, changes in beta and alpha cells in CF were largely distinct from those previously observed in T1D and T2D, which supports that CFRD is a distinct form of diabetes. This is likely due in part to different signals that endocrine cells receive in each form of diabetes where, in CF, they receive matrix interaction, inflammatory and proliferative signals primarily from areas of tissue damage. By comparison, beta cells in T1D respond to immune infiltration and up-regulate different inflammatory processes such as interferon signaling^72^ and in T2D respond to elevated glucose and lipid levels leading to glucolipotoxicity^73^. We observed an overall increase in processes and networks related to insulin secretion in beta cells in CF indicating potential compensation for persistent hyperglycemia, likely due to multiple factors including reduced overall beta cell mass, barriers to insulin release into the blood, and insulin resistance, which may further contribute to beta cell dysfunction and loss. In this way, beta cells in CF may have some similarities to T2D pathogenesis, where in the latter compensation primarily occurs due to insulin resistance, and which may also help explain the sharing of genetic risk factors for CFRD with T2D likely involved in insulin secretion^66^.

Previous studies of the pancreas in CF have found increased numbers of lymphocytes and activated stellate cells^22,23,39^, and higher levels of interleukin 1 beta^31^, which we confirmed in our data. While we did not find evidence for an increase in T cells as reported by those studies, our study had limited resolution of specific T cell sub-types, potentially due to lower transcript abundance in T cells, and thus alternative approaches like highly multiplexed immunostaining may be required to better understand immune-related changes in CF. We observed increased pro-fibrotic activity in stellate cells and inflammatory activity in immune cells, and our analyses revealed that crosstalk between these cell types as well as localization to areas of tissue damage may be responsible for driving increases in activity. Stellate, immune and endothelial cells were all in closer proximity in the pancreas in CF which may help explain the increased degree of signaling and activation; for example, macrophages received canonical recruitment signals such as CXCL, MIF and CSF from both activated stellate cells and endothelial cells, as well as collagen signaling from stellate cells which associated with upregulation of pro-fibrotic and growth-related processes.

There are several limitations of our study that future work can build on in multiple ways. Almost all CF donors in our study were between the ages of 20 and 30, and thus we had limited ability to define changes in the pancreas across different age groups particularly in very young individuals. Studies of the pancreas in CF have found increased activity of immune and stellate cells from very early ages^39^, supported by the dramatic increase in immune cells observed in a two-week old individual with CF profiled in our study. Pancreatic damage in CF begins very early in life, and understanding the initial sources and timing of pancreatic dysfunction is essential to preventing later onset of CFRD. In addition, as we analyzed all CF donors together due to the relatively limited sample size, increasing number of profiled donors will enable characterizing differences in CF stratified by variables such as CFRD or *CFTR* mutation status, which in the former case can reveal factors that either promote or protect against the development of diabetes in the context of CF. Furthermore, all CF donors in our study were collected prior to the availability of highly effective CFTR modulator therapies. While this provides critical baseline data regarding the pathology of the CF pancreas in the absence of HEMT, determining the impact of these therapies on pancreatic remodeling and cellular function will be important for understanding the current landscape of diabetes pathogenesis in CF. Finally, more detailed interrogation of cell types and sub-types across modalities such as spatial proteomics will enable deeper resolution and insights into the CF pancreas.

Overall, comprehensive single cell and spatial profiling of the human pancreas in CF revealed substantial new insights into cellular changes in the pancreas CFRD and provided new in-roads to preserving beta cell mass and function in CF.

## Methods

### Sample collection

De-identified human pancreas samples were obtained from several sources, based on the limited availability of CF samples. Donor pancreas samples were obtained from the Network for Pancreatic Organ Donors with Diabetes (nPOD; RRID:SCR_014641): n=5 donors with CF, n=5 donors with no-CF and no diabetes as controls. Deidentified autopsy pancreas samples were received from the National Disease Research Interchange (NDRI; RRID:SCR_000550; n=2 donors with CF, n=2 control) and the Pathology Department at Seattle Children’s Hospital (SCH; n=3 donors with CF and 3 control) from consented patients. For all sources of tissue, control samples were selected to match sex- and age- to the CF/CFRD samples as closely as possible. Human donor tissue studies were approved by the IRB of the University of California San Deigo.

For multiome analysis, snap-frozen samples were obtained. One of the frozen CF samples from NDRI had an extremely long post-mortem recovery interval (754 hours), precluding isolation of RNA. As such this sample and its matched no-CF/ND control sample were excluded from analysis. For spatial (CosMx profiling) sections from formalin-fixed paraffin embedded tissue were obtained. For spatial profiling we found that only donor samples, or autopsy samples with relatively short term (<12 year) storage at ambient temperature, exhibited sufficient RNA signal for analysis (described in more detail below), precluding the use of samples from a larger collection of CF/non-CF donors^31^.

### Nuclei Isolation from Frozen Pancreatic Tissue for Single cell multiome

Nuclei were isolated from frozen pancreas tissues of variable mass using Dounce homogenization. All procedures were performed on ice, and all reagents were pre-chilled. Tissues were lysed in NIM-DP-L buffer (0.25 M sucrose [Sigma S1888], 25 mM KCl [Invitrogen AM9640G], 5 mM MgCl₂ [Invitrogen AM9530G], 10 mM Tris-HCl, pH 7.5 [Invitrogen 15567-027], 1 mM DTT [Sigma D9779], 0.1% Triton X-100 [Sigma-Aldrich T8787], 1× Roche Complete Protease Inhibitor Cocktail [Roche 5056489001], and 1 U/μL RNasin Ribonuclease Inhibitor [Promega N211B]).

For homogenization, 10–15 mL of NIM-DP-L buffer was used per sample. Tissue disruption was performed using a loose-fitting pestle (15–20 strokes) until no visible tissue fragments remained, followed by an additional 10–15 strokes with a tight-fitting pestle. The resulting homogenate was filtered through a 30 μm cell strainer and centrifuged at 4°C, 10,000 rpm for 10 minutes. Supernatants were discarded, and the nuclear pellets were resuspended in sort buffer (1% BSA [Lampire Biological Laboratories 7500804] in PBS). Nuclei were stained with 2 μM 7-AAD (Invitrogen A1310) for 30 minutes.

After staining, nuclei were washed twice with sort buffer and sorted via fluorescence-activated cell sorting (FACS). Post-sort, library concentration was quantified using the Qubit dsDNA HS Assay Kit (Thermo Fisher Scientific) and fragment size distribution assessed on a TapeStation High Sensitivity D1000 ScreenTape (Agilent). Libraries were sequenced on both NextSeq 500 and NovaSeq X Plus platforms (Illumina).

### 10x Multiome Pilot Experiment

To ensure tissue processing and library preparations were successful on highly remodeled samples from donors with CF, we first conducted a small pilot experiment on nuclei from 5 donors from NDRI, CF (n=2) and non-CF (n=3). Nuclei from each donor were permeabilized by incubation on ice for 2 minutes in permeabilization buffer containing: 10 mM Tris-HCl (pH 7.5) [15567-027], 10 mM NaCl [AM9760G], 3 mM MgCl₂ [AM9530G], 0.01% IGEPAL CA-630 [Sigma I8896], 0.01% Tween-20 [Sigma P7949], 0.001% digitonin [Promega G9441], 1 U/μL RNasin [N2515], and 1× Roche Complete, EDTA-free Protease Inhibitor Cocktail [5056489001]. Reactions were quenched by addition of wash buffer (1% fatty acid-free BSA [7500804], 10 mM Tris-HCl pH 7.5, 1 mM DTT, 3 mM MgCl₂, 10 mM NaCl, and 0.01% Tween-20). The permeabilized nuclei were passed through a 40 μm Flowmi cell strainer (Bel-Art) to remove aggregates and resuspended in 1× Nuclei Buffer (provided by 10x Genomics). Nuclei concentrations were determined using a hemocytometer. Two of these initial five samples had very low-quality data after sequencing and were not used to construct our final Multiome map or run any analyses.

### 10x Multiome Primary Pooling Experiment

For the initial pooling experiment, nuclei from non-CF donors (n = 9) and individuals with CF or CFRD (n = 9) were processed and sorted independently using the same nuclei sorting and permeabilization methods as in the pilot experiment. To ensure equal representation across conditions, 122,400 nuclei from each pool (non-CF and CF/CFRD) were combined into a single suspension. This pooled sample was then distributed into eight reactions at ∼30,000 nuclei per lane on the Chromium Controller for downstream processing using the Chromium Next GEM Single Cell Multiome ATAC + Gene Expression Kit (v2, Rev F).

### 10x Multiome Cell Enrichment Pooling Experiment

Based on transcriptional profiling and quality metrics from the primary dataset, donors were re-stratified into three groups based on nuclear yield: Low CF/CFRD (n = 6), High CF/CFRD (n = 3), and Control (n = 8). All samples were processed through nuclei sorting and permeabilization as in the pilot experiment. For targeted enrichment, nuclei pools were combined at a 2:1:1 ratio (Low:High:Control) prior to loading onto the Chromium platform. A total of eight lanes were prepared with ∼30,000 nuclei per lane.

### Single cell multiome sequencing

Single-cell Multiome libraries were generated following all manufacturer instructions (Chromium Next GEM Single-cell Multiome ATAC + Gene Expression Reagent Bundle, 1000283; Chromium Next GEM Chip J Single cell, 1000234; Dual Index Kit TT Set A, 1000215; Single Index Kit N Set A, 1000212; 10x Genomics) with 7 cycles for ATAC index PCR, 7 cycles for cDNA amplification, 13-16 cycles for RNA index PCR. The final libraries were quantified using a Qubit fluorimeter (Life Technologies) and library size distribution was checked using Tapestation (High Sensitivity D1000, Agilent). Finally, all libraries were sequenced on NextSeq 500 and NovaSeqX Plus (Illumina) with read lengths (Read1+Index1+Index2+Read2): ATAC (NovaSeqX Plus) 50+8+24+50; ATAC (NextSeq 500 with custom recipe) 50+8+16+50; RNA (NextSeq 500, NovaSeqX Plus): 28+10+10+90.

### Sample genotyping and imputation

For all donors in the Multiome pools, we sampled 10-15mg of frozen tissue and extracted genomic DNA using the Monarch Genomic DNA Purification Kit (NEB #T3010L). Samples were next eluted in 70-100uL UltraPure Distilled Water (Invitrogen 10977-015), and then concentrated via speed vacuumed to a concentration of 50ng/uL if necessary. All DNA samples were genotyped with the Illumina InfiniumCoreExome-24v1-4_A1 array at the UCSD IGM Genomics Center.

First GenomeStudio v2.0.5 was used to cluster positions and extract SNP statistics. Then genotypes were converted to plink format by using the GenomeStudio plink input report plugin and the hg38 cluster file. Samples run on different arrays were merged using plink v1.9 and then variants were filtered for quality. We retained all variants with a minimum missing call rate of 0.05, minor allele frequency of at least .01, and Hardy-Weinberg equilibrium p-value greater than 1e-5. Finally we generated VCF and frequency files and prepared variants for imputation by using the HRC-1000G-check-bim.pl script from the McCarthy Group Tools (https://www.chg.ox.ac.uk/~wrayner/tools/, v4.3.0). Genotypes were imputed using the TOPMed r3 panel and the TOPMed Imputation Server^74^. Imputation was performed with Eagle (v2.4) using the Array Build Hg38 with phasing on and r2 filter off. After imputation, variants were filtered by imputation quality (r2>0.9 and minor allele frequency (MAF) > 0.01.

### CosMx spatial transcriptomics assays

For spatial profiling, benchwork was done at the Experimental Pathology shared resource (Fred Hutch Cancer Center, Seattle, WA). Formalin-fixed, paraffin embedded (FFPE) pancreas sections from CF/CFRD (n=9) and no-CF/ND (n=9) donors/autopsy cases from nPOD (n=6 per group) and from SCH (n=3 per group). CF/CFRD samples were from the same donors/cases as the multiome analysis with 1 additional CFRD nPOD donor (and no-CF/ND control) for which there was FFPE but no frozen pancreas available. No histological samples were available from NDRI. Prior to spatial profiling, *in situ* hybridization (RNAscope for *PPIB)* was performed on sections from all samples to assess RNA quality. From this analysis, a total of 6 CF/CFRD samples (4 from nPOD, 2 from SCH) and a corresponding 6 no-CF/ND (4 from nPOD, 2 from SCH) were selected for further analysis.

Fresh FFPE sections (4 µm) were cut from prioritized blocks for spatial profiling and deparaffinized slides had flow cells attached for CosMx Human RNA 1000-Plex assay (Nanostring, Bruker, Billericia, MA). Samples first underwent multiplexed immunofluorescence (Hs Universal kit – CD298+B2M + Hs PanCK/CD45 marker kit) to identify tissue features. Fields of view (FOV, selected by RLHM and the Experimental Pathology team) on CF/CFRD sections were selected to encompass most of the tissue due to low cellularity, while on no-CF/ND tissues, FOVs were selected to ensure good representation of islets and exocrine tissue. For both groups, FOVs were also prioritized for areas with good RNA quality (typically around the edge of a section). Multiple reporter sets containing antibodies linked to oligonucelotide tags were then cyclically applied to tissue sections with high-resolution spatial imaging completed for each reporter set. Following CosMx assay, sections were stained with H&E to permit analysis and registration of tissue landmarks to the spatial data.

### Multiome data quantification and alignment

Cell Ranger ARC (v2.0.2, 10x Genomics) was used to process all raw Multiome sequencing data. The count command was used to align the sequencing reads to the GRCh38 genome reference with GENCODE v32 annotations. Per-modality bam files and feature barcode count matrices were also generated.

### Processing and QC of Multiome counts data

We individually processed the data from each separate sequencing experiment. Duplicate reads were removed from the ATAC bam files and these were converted to the tagAlign format and used to quantify all ATAC counts in 5kb windows tiling the human genome with bedtools^75^. Background RNA contamination was cleaned from the RNA counts matrices using SoupX^76^ and the auto method for estimation of each cell’s contamination fraction (v1.6.2). All pooled samples were deconvoluted using demuxlet^77^ (v2) (https://github.com/statgen/popscle) with the VCF created from all imputed pool donor genotypes.

To identify barcodes representing single cells of high sequencing quality, we used the following per-barcode metrics: RNA counts >250, ATAC counts >1000, RNA mitochondrial reads <1%, ATAC Transcription Start Site enrichment (TSSe) > 1. Additionally, we removed all barcodes identified as cross-sample doublets by demuxlet, all barcodes identified as multiplets from Cell Ranger, and all barcodes identified as doublets by Amulet^78^ (v1.1).

### Multiome clustering and cell type annotation

To identify a select set of 5kb ATAC windows that distinguish cell types we first merged data from 5 sequencing lanes: all three samples sequenced individually from the pilot experiment and two lanes of pooled samples. All data passing QC filters from these experiments were clustered with the following protocol: RNA counts were normalized with SCTransform^79^, and then used to perform PCA. ATAC counts were normalized with term-frequency-inverse document frequency^80^ (TF-IDF), the FindTopFeatures command was used to identify the features with the most counts, and then singular value decomposition (SVD) was performed. We applied Harmony^81^ to the RNA and ATAC embeddings to correct for donor and sequencing lane if applicable. We then ran UMAP on each separately with the top 50 PCs, excluding PC1 from ATAC because this is usually associated with sequencing depth. Finally, the Weighted Nearest Neighbor^41^ algorithm was used to project cells in a low dimension space using both modalities and we reran UMAP and used the Leiden algorithm^82^ (resolution=0.25) to cluster barcodes. Finally, the FindVariableFeatures command was run on the ATAC data to identify the top 50,000 most variable 5kb windows across samples.

To merge all donor and pool samples into one object, we subset the ATAC 5kb matrices to the previously identified 50,000 variable windows and then reperformed the same clustering procedure. After this there were a few very small clusters with no strong marker gene expression and lower than average RNA counts. We removed these from the map, as they were likely doublets or low-quality cells, and then reperformed the clustering procedure. The cell type identities of all clusters in the final map were manually annotated using known islet cell type marker genes. For select clusters which represented subtypes of known cell types, we used Seurat’s FindAllMarkers() command and fGSEA^83^ to identify genes and pathways enriched in each and used these to annotate each.

### Peak calling on snATAC-seq data

Peaks were called on all final cell types using the callpeak tool from MACS2^84^ (v2.2.7.1), merged tagAlign files with all reads for all cells in the cell type, an FDR cutoff of 0.05, and the –nomodel and –keep-dup all flags. To identify a single condensed peaks set for downstream analyses we first limited all peaks to 300bp around the peak center. We then created clusters of peaks based on overlap and for each we assigned the peak with the highest read count at the summit as the reference peak. We repeated this process on all peaks that did not overlap any reference peaks until no peaks remained. Finally, we filtered this set to only peaks with a MACS2 -log10(q-value) > 10. For each cell type, we identified which peaks in this final set were accessible in the cell type by intersecting each cell type’s original peak calls with the final peak set with bedtools intersect.

### Construction of cell type gene regulatory networks with SCENIC+

We used SCENIC+^42^ to identify TF-cRE-gene regulatory networks, or eRegulons for all cell types with consistent representation across donors. First, we subset the paired snRNA and snATAC map to 50,000 representative cells using Seurat’s sketching feature (v5.2.1)^85^ and the “LeverageScore” method. To prepare inputs for SCENIC+, pycisTopic (v2.0a0) was used to perform topic modeling of the final peak set, identifying 45 topics. Next, all peaks were binarized and assigned to topics with the ‘otsu’ method and the ‘ntop’ method. Peaks differentially accessible between cell types were identified with the find_highly_variable_features() and find_diff_features() commands and default parameters from pycisTopic. Next, cluster-buster was used to create a custom cistarget database using the hg38 genome and the final peaks set, with 1kb region padding. Finally, SCENIC+ was run using the snakemake implementation (v1.0a2), the chromatin inputs made with pycisTopic and cluster-buster, and the per-cell RNA counts matrix.

Pathway files in were constructed for all TF modules using all SCENIC+ predicted target genes for each TF. To identify TF modules enriched in specific cell types, per-cell gene-based AUC values were calculated for all cells in the full map using the score_eRegulons() function. Then all AUC values were binarized so that cells with an AUC value more than 1 standard deviation above the mean were considered to have an active TF module and all others were inactive. TF modules that were active in more than 40% of the cells in a cell type were classified as cell type enriched.

To identify pathways enriched in target genes of each TF in a cell type and disease specific manner, we gathered all predicted target genes for a TF and filtered them to all genes with a per-donor average TPM>10 in the relevant cell type and disease group. We next removed any ribosomal genes, mitochondrial genes, or unannotated genes before running enrichR^86^ pathway enrichment on each set of genes.

### Identification of cell type-specific marker genes, cREs and motifs

To determine marker genes and cREs for each cell type, per-donor pseudobulk matrices were generated from all cells in the cell type and all cells in other cell types. For RNA analyses, all genes were used and for ATAC, all peaks in the final set were used. DESeq (v1.46.0) was ran using a formula to predict the cell type identity from gene expression, including sample ID as a covariate. For each cell type only control (no-ND/no-CF) donors were used and only if the donor had more than 20 cells in the cell type of interest. All cell types with less than 2 donors passing this filter were skipped. HOMER^87^ (v5.1) was used to identify TF motifs enriched in the cell type-specific peaks identified by DESeq. For each cell type we split significant marker cREs at FDR < 0.1 based on change in direction. For each group of cell type specific, directionally altered cREs associated with a cell type, we conducted motif enrichment analysis using HOMER’s findMotifsGenome.pl. As background, we used all peaks identified within the corresponding cell type, applying the default window size of 200 bp.

### Tests for cell types with altered abundance in CF pancreas

To robustly compare the proportion of cell types between donors with and without CF, we first extracted and square root scaled the proportion of each cell type per donor within its relative compartment (exocrine, endocrine, stellate, immune, other). Then we performed a two-sided Welch’s t-test to compare the mean proportions between the two donor groups and used Benjamini-Hochberg multiple test correction to identify significantly altered cell type proportions. All tests were performed separately using proportions from the Multiome data and CosMx data.

### Tests for genes, pathways and TF modules differentially expressed with CF status

To identify genes changing with CF status in different cell types, DESeq2^88^ was run on per-donor cell type pseudobulk total RNA counts matrices. Only counts from donors with at least 20 cells in the cell type of interest were included in these analyses, and we only tested genes with at least 100 counts total. Analyses were run on all cell types that had at least two donors in each test group (CF and control). DESeq was run with the following formula: CF status ∼ age + BMI + percent mitochondrial counts + tissue source, where all continuous per-donor variables were scaled first. To identify pathways altered in CF, we first selected all genes with FDR<0.1 differences in gene expression from DESeq and split them into two groups based on the direction of effect. We next removed any ribosomal genes, mitochondrial genes, or unannotated genes before running enrichR^86^ pathway enrichment on each set of genes. Finally, fGSEA was also run on the DESeq results to identify TF modules that were up or down regulated with CF status. The same set of genes were removed before these analyses, in addition to genes in the top 20 markers of any cell type, to remove the effects of contamination from over-represented cell types. A rank value was then calculated for all remaining genes, comprised of the gene’s log2FoldChange and -log10(p-value) from DESeq. The rank values were used to run fGSEA on a custom set of pathways comprised of all genes predicted to be regulated by a TF by SCENIC+ (TF-gene correlation > 0.2), for each TF with TPM > 10 in the cell type.

### Tests for peaks and motifs differentially accessible with CF status

We used a very similar approach to identify differentially accessible ATAC peaks in CF. First, per donor pseudobulk ATAC counts matrices were generated using our final peak set. The same per-donor filters were used to identify which donors and cell types could be tested confidently. Additionally, only peaks with more than 100 total counts and which overlapped a peak initially found in per-cell type peak calling were tested. DESeq was run with the same formula, however TSS enrichment was used instead of percent mitochondrial counts.

Transcription factor motif variability was analyzed using chromVAR^89^ (v1.24.0). Accessible chromatin counts per million were generated using 300 bp union peaks, and peak coordinates were used to construct GRanges objects. Motif binding was predicted using motifmatchr (v1.24.0) with BSgenome.Hsapiens.UCSC.hg38 as the reference for motifs from the JASPAR 2022 database. To account for GC-content bias, we applied chromVAR’s addGCBias function. Motif deviations were then calculated using computeDeviations. The resulting motif x barcode deviation scores were pseudobulked into motif x donor deviation scores by averaging across cells from each donor. Finally, to identify differentially accessible motifs associated with disease, we applied linear models to the pseudobulked chromVAR deviation score matrices for CF vs. Control donors, using the base lm function in R (v4.3.3) and the formula: motif∼ cf_status + gender + age.scaled + BMI.scaled + tissue_source + avg_TSSe.

### Construction of a ductal cell trajectory

To identify cells potentially transitioning between conventional ductal and high-mucin ductal identities, we used Monocle3^90^ (v1.3.4) to perform trajectory analysis. First, we extracted the raw gene expression counts and UMAP coordinates for all ductal and high-mucin ductal cells from donors with CF. Then Monocle’s learn_graph() method was used to identify a trajectory in these cells with the following parameters: ‘annoy’ nearest neighbor method, Euclidean nearest neighbor metric and a maximum distance ratio of 1, a max of 10 trees with minimal branch length of 20, 200 centers and a closed loop. Next, we selected the node within the ductal cell cluster that was the furthest away from high-mucin ductal cells to calculate pseudotime from. Finally, to identify features changing with cell identify, we selected all cells with a pseudotime between 5 and 10, as this was the range in which proportions of the two cell types changed the most, and tested the Spearman correlation of gene expression, peak accessibility, and TF module AUC with pseudotime. The cor.test() function was used to calculate a two sided p-value associated with this correlation and a smoothed representation of each feature over pseudotime was calculated with a rolling mean over 2,000 cells for visualizations.

### Processing and QC of CosMx data

All CosMx data were processed through CosMx pipelines via the AtoMx Spatial Informatics Platform which identifies cell boundaries with CellPose and generates counts matrices for all cells. First, to identify cells which were likely properly segmented we filtered for all cells with mean DAPI and membrane staining signal greater than or equal to 100 and cell area greater than or equal to 25µm^2^ and less than or equal to 200µm^2^. Additionally, cells at the 36um regions bordering all FOVs were also excluded as they likely are subject to distortion from the Barrel effect^91^. Next, cells with high quality RNA data were selected with the following per-cell filters: greater than or equal to 250 RNA counts, greater than equal to 50 unique genes, and less than 3 counts from negative probes. Due to differences in RNA quality previously observed in highly remodeled CF samples, we used slightly adjusted filters for all cells from CF donors: greater than or equal to 75 RNA counts, greater than equal to 25 unique genes, and less than 3 counts from negative probes.

### Generation of an annotated single cell map from CosMx data

Scanpy^92^ (v1.10.3) was used for all clustering of spatial transcriptomics data. First, gene counts were normalized to 10,000 counts per cell with the normalize_total() function including highly expressed genes. Counts were log transformed, then used to perform PCA to generate 50 PCs which were corrected for sample differences with harmony integration^81^. Finally, scanpy’s neighbors() function was used, a UMAP representation was calculated with min_dist of 0.3 and spread of 1, and finally clusters were identified with the leiden algorithm and a resolution of 1.

An initial cell type annotation was completed manually using the expression levels of known cell type marker genes in the leiden clusters. However, it was challenging to confidently distinguish rare cell types including stellate and immune cells with this method. Thus, to identify fine-grained annotations of all cell types in the multiome map, we used scANVI^45^ from scvi-tools (v1.2.0) to perform label transfer of these annotations onto the spatial map. First, both gene expression matrices were cut down to only genes which were found in both datasets, excluding the INS gene which is often highly represented in contamination profiles, for a total of 964 genes. The multiome data was used to create an scVI model with default scArches parameters. This model was then trained to identify the known cell type labels for a maximum of 20 epochs, using 100 samples per label. To decrease training time, a GPU node was used for model training via PyTorch. Next, the CosMx counts matrix was used to create a second query model that was trained on the reference model for a maximum of 100 epochs, with 0 weight decay. Cell type labels were then predicted with the predict() function. In order to more accurately separate endocrine cells, this process was then repeated just on all cells identified as endocrine cells (beta, alpha, delta, gamma or epsilon), using a reference model with only endocrine cells.

### Identification of tissue niches with BANKSY

To identify conserved tissue domains across all spatial samples, we first arranged all samples into the same coordinate system by staggered all x-axis coordinates, so all samples were separated by 1.5 times the max width of any sample. Regions of contiguous FOV were identified for each sample and annotated as tissue regions. For one sample (SCH_CF2), we found strong batch effects along a vertical axis in a contiguous region and divided this into two regions. Standard BANKSY^46^ preprocessing was applied using the normalize_total() and filter_hvg() commands for 2,000 variable genes using the Seurat method. Next, spatial weights were calculated based on distance with the initialize_banksy() command using 15 neighbors, the Azimuthal Gabor filter, and gaussian decay. BANKSY matrices were generated using a lambda value of 0.9, to prioritize the expression profiles of a cell’s neighbors above its own profile. PCA was then run and the 20 PCAs generated were corrected using harmony for the sample identify, sample region, and donor tissue source. Finally, UMAP was run on the harmony corrected PCs and niches were identified using the leiden algorithm with resolution=0.25. This process identified a total of 11 niches, however two niches were comprised of less than 1,000 cells and not found in all samples, so we did not consider them for any downstream analyses. All BANKSY commands were from the python implementation of the package, Banksy_py (https://github.com/prabhakarlab/Banksy_py).

### Immunofluorescence

4µm sections of formalin-fixed, paraffin-embedded (FFPE) pancreas samples (consecutive sections to those used for CosMx) were subjected to a standard rehydration process with 5 minute washing in each reagent in the following order: xylene x 3, 100% ethanol, 95% (v/v) ethanol, 70% (v/v) ethanol, and 10 minutes in ddH2O. Samples were incubated in 0.3% H_2_O_2_ in methanol for 20 minutes to minimize background autofluorescence. Heat-mediated antigen retrieval was performed (EDTA buffer, pH 9) for 20 minutes at 95℃ followed by permeabilization (1% Triton X-100 in PBS, 5 minutes at room temperature), washing (PBS 3x5 minutes) and blocking (2% (v/v)normal donkey serum, 1% w/v bovine serum albumin, 0.01% w/v sodium azide; 0.2% (v/v) Triton X-100 in PBS) for 30 minutes. A primary antibody solution was prepared (antibodies listed in **Supplementary Table 23**) in blocking buffer, and incubated overnight at 4℃ in a humidified chamber. Following washes (PBS 3x5 minutes with gentle rocking), a secondary antibody solution (antibodies listed in **Supplementary Table 23**) in blocking buffer and incubated for 1 hour at room temperature in a humidified chamber. Samples were washed PBS 3x5 minutes with gentle rocking), followed by incubation with TrueView Quenching Kit (vendor: Vector Labs, SP-8400-15) according to the manufacturer’s instructions and final wash (PBS 1x5 min). Sections were counterstained with DAPI (1 µg/ml; Life Technologies, 62248, 10 minutes at room temperature), washed (PBS 3x5 minutes with gentle rocking), coverslipped with Prolong Glass mounting media (ThermoFisher, P36980) and cured overnight at 4℃.

Whole tissue sections were imaged with Leica Thunder deconvolution widefield fluorescence microscope (Leica Microsystems) using a 20x Plan Apochromat air objective lens (numerical aperture 0.8) at room temperature. Tiled images were acquired with Leica LASX software v5.3.0. Subsequently Leica Thunder Computational Clearing (CC) was applied.

### Calculation of cell-cell proximity values

The co-occurrence method implemented in the TACCO package^93^ (v0.4.0.post1) was used to calculate the likelihood of cells to be spatially proximal at increasing distance thresholds. For each sample, the co_occurrence() function was used on all pairs of cell types with more than 50 cells in the sample, at 25µm distance bins between 0 and 2,000µm using a sparse matrix representation. Due to memory constraints two very large samples were split into multiple regions before running and all cell-cell proximity values were averaged across regions for each distance bin. nPOD_CTL5 was split into its 3 contiguous FOV tissue regions and SCH_CTL5 was split into two regions lengthwise. TACCO automatically scales all proximity values by the composition of the 15 closest cells and did not require normalization for differences in tissue density between samples. To compare changes in cell-cell proximity between control and CF samples, for each pair of cell types the average probability of proximity was calculated from all bins between 0 and 50µm.

### Identification of cell-cell signaling interactions changing with CF status

To predict cell-cell interactions in a spatially informed manner, CellChat^94^ was run separately on each spatial sample. All cell distances were calculated in microns using a conversion factor of 0.12028 (CosMx standard) and the spot size was set to the minimum cell-cell distance in microns. Next, a CellChat object was created using all cells and all genes overlapping between our 1,000 gene panel and the CellChatDB human database. Subsequent analyses were performed on a total of 341 ligand or receptor genes which were highly expressed and variable. The expression matrix was subset to over expressed genes and interactions and then communication probabilities were computed using spatial information. The following parameters were used in the computerCommunProb() function: the “truncatedMean” was used with a trim of 0.1, population size and distance were used, with a max interaction range of 250µm and a distance scale factor of 0.1, and all contact dependent interactions were calculated with a contact range of 10µm. For one very large sample (SCH_CTL5) this process was run separately on two pieces of the tissue, split lengthwise and all resulting communication probabilities and p-values were averaged.

To test for cell-cell signaling interactions that were different in control and CF donors, we first computed a weighted probability value for every interaction. First all CellChat p-values were adjusted for multiple tests with the Benjamini Hochberg method, and then only ligand-receptor pairs that were significant (FDR<0.1) in at least one pair of cell types in at least one sample were retained. Weighted probability values were calculated by setting all p-values of 0 to 0.01 and then multiplying the probability by the -log10 of this p-value. We found that the number of FDR significant interactions identified per sample was strongly correlated to the number of cells in the sample. Thus, per-sample pathway interaction values were computed as the sum of all weighted probability values for all ligand-receptor pairs in the pathway and then scaled by the total number of cells in that sample. A similar process was applied to calculate sample values of total or specific signals sent by or received by a cell type. In all cases the average weighted probability was scaled by the number of cells per the cell type of interest in each sample. Finally, for comparisons of interactions between two specific cell types, either for all interactions or a specific pathway, all average weighted probability values were scaled by the number of cells in the receiver cell type. All comparisons between control and CF sample values were performed with a two-sided Welch’s t-test.

### Receptor expression correlation analyses

We first calculate per-cell pathway enrichment values using UCell and the multiome data. Next, for a cell type and signaling pathway of interest, per-donor pseudobulk values were calculated for all unique receptors in the pathway (mean CPM), all genes with an average TPM > 10, UCell pathway enrichment (mean UCell score), and SCENIC+ TF modules (mean AUC). Then the Spearman correlation was calculated between all receptors with a mean TPM > 1 and SD > 1 in the cell type and all genes (average TPM >10), pathways (UCell score SD > 0.01), and TF modules (TF TPM > 10 and AUC SD > 0.01). Finally, multiple test correcting was performed using the Benjamini-Hochberg method to identify significant correlations.

### Annotation of tissue structures in H&E images

A board-certified pathologist (GHD) performed manual annotation for regions in 10 of the 12 spatial samples, using the H&E images. All regions contiguous with selected FOVs were annotated from each tissue. The following classes of tissue features were annotated in more than half of the samples: adipose tissue, artery, collagen, dense collagen, nerve, vein, and vessel. For both adipose tissue and vessels there was a large range in feature area and thus we split these into subclasses based on the following thresholds: small adipose tissue (<=100,00µm^2^), large adipose tissue (>100,000µm^2^), small vessel (<=700µm^2^), medium vessel (>700µm^2^, <=8,000µm^2^), and large vessel (>8,000µm^2^). Annotations were exported from QuPath with a custom groovy script that collected all points in the perimeter of each annotation. These were then converted to microns using the sample specific down sampling number from QuPath export (between 4 and 8) and 0.4965 as the pixel size unit.

### Alignment of H&E images with CosMx cell coordinates

To align CosMx cell coordinates with the coordinate system of the H&E images, we used the STalign^95^ tool (v1.0). First, we separated each sample into regions of contiguous FOVs and gathered all cell center coordinates for each region. Normal STalign pre-processing steps were applied to the H&E image including normalization, transposing and rasterization (dx=100) with default parameters. Then for each region, between 3 and 7 landmark points were identified manually to perform an initial affine transformation on the CosMx coordinates. Following this, STalign LDDMM was run with all default parameters using these coordinates and the rasterized H&E image. Finally, the coordinates returned by this were multiplied by the downsample term used to export each H&E image from QuPath (ranged from 4 to 8) and the QuPath pixel size (0.4965 for all samples).

### Calculation of cell type changes with proximity to tissue structures

All annotation object coordinates were imported to python as shapely.Polygon objects (v2.0.7)^96^ and for each annotation class an STRtree was constructed to hold all annotations per sample. Next, all cells in the annotated regions of the sample were converted to shapely.Point objects and compared to the STRtree to calculate the minimum distance from each cell to an object of the annotation class. This was repeated for all annotated samples and all annotation classes.

To identify cell type populations enriched or depleted with proximity to tissue structures, the proportions of all cell types were calculated at radial distance bins around each annotation class. This was performed separately on all cells from control samples and CF samples in 5µm bins from 0 to 4,000µm and for all tissue structure annotation classes. For each annotation class and disease group, sparse distance bins were identified as the first 5µm bin over 500µm with less than 20 total cells. All subsequent calculations were only performed up to the first sparse distance bin. Next, the Pearson correlation of each square root-scaled cell type proportion and the distance bin was calculated for each annotation class and disease. The cor.test() function from R was used to perform these correlations and generate a p-value for significance based on a t-distribution with n-2 degrees of freedom. Finally, all p-values were corrected for multiple testing with the Benjamini-Hochberg method.

To identify genes with per-cell type expression changing with spatial proximity to tissue structures, we performed similar analyses. First, for each cell type and disease group, we identified the 50% more variable expressed genes by first selecting genes with more than 0 counts in at least 5% of the cells in the cell type and then taking the half with the highest standard deviation. Next, we calculated the Pearson correlation between per-cell gene counts and the minimum distance to a structure within the annotation group for each of these genes and annotation class. All resulting significance values were corrected for multiple testing with the Benjamini-Hochberg method. Finally, to identify cellular trends in genes correlated to distance from a tissue structure, we used enrichR^86^ to test for pathways enriched in all genes with an FDR<0.1 distance correlation and split by correlation direction using the GO_Molecular_Function_2021 and Reactome_Pathways_2024 databases.

### Classification of cell type defined tissue structures

To identify spatial regions likely to represent large pancreas ducts, the DBSCAN^97^ clustering algorithm was applied to the spatial coordinates of all high-mucin ductal cells. For each cell, the cell center x- and y-coordinates were extracted and then DBSCAN was run on them with a max distance of 75µm and a minimum number of neighboring cells of 10 for all center points. For one sample, nPOD_CTL6, which had very small FOV regions a max distance of 50µm was used to define large ducts. After cells belonging to separate ducts were identified, the concave hull for each separate duct was calculated as follows: all x and y points from cell coordinates were min-max scaled and then an alphashape object (v1.3.1)^98^ was created with these. The alpha value used for this was based on the number of cells in the duct, alpha=30 if the duct had more than 500 cells, alpha=2 if the duct had less than 100 cells, otherwise alpha=15. After this shape was created, it was denormalized back to the original coordinates. All potential radii for each duct were calculated by computing the Euclidean distance from each boundary point to the duct centroid.

The same method was applied to all spatial coordinates of endocrine cells (beta, alpha, delta, gamma, epsilon) to define individual pancreatic islets. For most samples, DBSCAN was run with a maximum distance of 54.126µm (450px) and a minimum of 30 neighboring cells per center. However, for three extremely fat infiltrated CF samples with small islets (nPOD_CF1, nPOD_CF4, nPOD_CF6), the minimum neighbors per center was reduced to 10. The same procedure was used to create an initial concave hull for each islet with alphashape. Then, a 1.2028µm(10px) buffer was added to the perimeter of each the denormalized shape and all separate polygons that were within 0.6014µm (5px) of each other were merged. This allowed for finer separation of islets that DBSCAN couldn’t separate. Next, all non-endocrine and non-exocrine cells found in each islet’s concave hull were also assigned to the islet. Seven outlier islets with extremely high numbers of cells assigned to them (3 standard deviation above the total mean) were removed. Finally, any cells initially assigned to multiple islets were assigned to the islet with the nearest centroid coordinates.

For each islet identified by the above procedure, the following metadata was gathered. First, all cells directly bordering the islet concave hull were collected by creating a cKDTree (scipy.spatial v1.15.2) from all non-endocrine cells and then performing query_ball_point() using the tree and all islet coordinates and a radius of 12µm. The proportions of cell types within and directly bordering the islet were calculated and square root scaled. The area of each islet was calculated by summing the areas for each cell using the Area.um2 output from AtoMx and density was calculated by dividing the area by the total number of cells per islet. The proximity to the nearest islet was calculated using the centroid points of each islet and minimum distance to each class of tissue structure annotations was calculated with the same method as used previously and using the islet centroid coordinates rather than cell coordinates. Additionally, for each cell within an islet the radial distance from the islet center was calculated using the centroid coordinates from the islet concave hull and the cell’s centroid coordinates. These values were min-maxed scaled to be directly comparable between islets of different sizes, and islet regions were defined off these normalized values as follows: core (≤0.4), middle (>0.4 and ≤0.75), and edge (>0.75). Cell type proportions within each region of every islet were also calculated. Finally, islets were classified based off their proximity to dense collagen (close if less than 50µm away) and large regions of adipose tissue (close if less than 100µm away).

### Identification of islet features and gene expression changing with disease

First, all continuous variables from the per-islet metadata collected were log2 scaled and the per-donor average value was calculated for each. Differences between control and disease islets were tested for each per-islet metadata feature using a Welch’s two-sample t-test on the per-sample averages (n=11) to control for large differences in number of islets per sample. Differences between islets classified on proximity to dense collagen and adipose tissue were calculated using an ANOVA test on all individual islet measures per classification category.

Additionally, tests were performed to identify cell type gene expression changing with various islet features. First all per-islet metadata values besides the number of cells (which is strongly correlated with islet area) were scaled between 0 and 1. Then, for the 50% most variable expression genes for each cell type we calculated the Pearson correlation of the gene’s expression and each islet feature and corrected resulting p-values with the Benjamini Hochberg method. Pathways enriched in genes significantly correlated or anti-correlated (FDR<0.1) with islet features, and that weren’t among the top 20 markers of any cell type, were identified using enrichR and the GO_Molecular_Function_2021 and Reactome_Pathways_2024 databases.

### Connectivity analysis of gene sets

We defined gene sets for connectivity mapping from two sources. First, we identified genes with significant up- and down-regulated expression in beta and alpha cells in T1D and T2D from a prior study^44^. Second, we identified genes with up- and down-regulated expression after CRISPR knockout of 5,049 genes from the Library of Integrated Network-Based Cellular Signatures (LINCS) project^99^.

### Comparison of genetic risk loci effects

To evaluate whether genetic risk loci for T1D and T2D show concordant effects in CFRD, we identified SNPs shared between each disease’s lead GWAS variants and the CFRD GWAS summary statistics. Lead SNPs from a T1D GWAS^100^ and T2D GWAS^101^ were intersected with all SNPs present in a CFRD GWAS^66^ using the base R intersect() function (R v4.1.1). Effect sizes (β) for each overlapping SNP set were then extracted, and Pearson correlation coefficients were computed using the base R cor() function to assess concordance between CFRD and T1D/T2D genetic effects.

### Pseudobulk comparisons of CFRD donors with Type 1 and Type 2 Diabetes donors

Raw read counts from pseudobulked RNA-seq data were merged across all donors from this project and donors with either type 1 diabetes (nPOD cohort^35^) or type 2 diabetes (City of Hope cohort^102^) for each cell type. A DESeq2 dataset (DESeq2 v1.34.0) was constructed using the design formula raw pseudobulked counts∼ gender + age + BMI + diabetes_status to model expression differences across samples. DESeq2’s vst function was applied to normalize the count data. To account for study-specific effects, batch correction was performed using the ComBat function from the sva package (v3.42.0), with the model normalized count matrix ∼ age + gender + BMI + diabetes_status and study as the batch variable. Principal component analysis was then conducted using the prcomp function in base R (v4.1.1) on the top 2,000 most variable genes per cell type to generate sample embeddings and gene loadings.

## Author contributions

H.M.M. performed single cell analyses and wrote the manuscript. S.C. contributed to single cell analyses. J.L. performed single cell assays. R.L.M. contributed to single cell analyses and created tools and visualizations for single cell data. M.P. contributed to single cell analyses. V.C.J. performed immunofluorescence experiments. G.H.D. contributed pancreas autopsy samples and annotated H&E images. A.W. and A.D-C. contributed to the generation of single cell assays. R.L.H-M. conceived the study, identified and coordinated samples and obtaining funding. K.J.G. conceived the study, supervised the study, obtained funding, contributed analyses, and wrote the manuscript.

## Data availability

Sequencing data will be deposited in GEO upon publication. Imaging data and count matrices will be deposited at BioImage Archive upon publication. Processed single cell objects and associated resources such as gene regulatory programs and accessible chromatin peaks will be made available at http://cfrdgenomics.org/. All other data and results are provided with the manuscript or available from the authors on request.

## Code availability

Code used in our study is deposited in the GitHub repository at https://github.com/Gaulton-Lab/cf-panc-multiomics.

## Acknowledgements

This research was performed with the support of the Network for Pancreatic Organ donors with Diabetes (nPOD; RRID:SCR_014641), a collaborative type 1 diabetes research project supported by Breakthrough T1D and The Leona M. & Harry B. Helmsley Charitable Trust (Grant#3-SRA-2023-1417-S-B). The content and views expressed are the responsibility of the authors and do not necessarily reflect the official view of nPOD. Organ Procurement Organizations (OPO) partnering with nPOD to provide research resources are listed at https://npod.org/for-partners/npod-partners/. We acknowledge the use of tissues procured by the National Disease Research Interchange (NDRI; RRID:SCR_000550). This work was supported by Cystic Fibrosis Foundation Award HULL22G0-CFRD to R.L.H-M, DartCF pilot award P30DK117469 to K.J.G, NIH award R01DK137209 to R.L.H-M, and CIHR award CERC-22-0023 to R.L.H-M. H.M.M. was supported by the DT O’Connor Scholar in Genetics and M.P. was supported by T32GM139790.

