## Supplemental figures for "Single cell and spatial characterization of the human pancreas reveals drivers of beta cell dysfunction in cystic fibrosis"

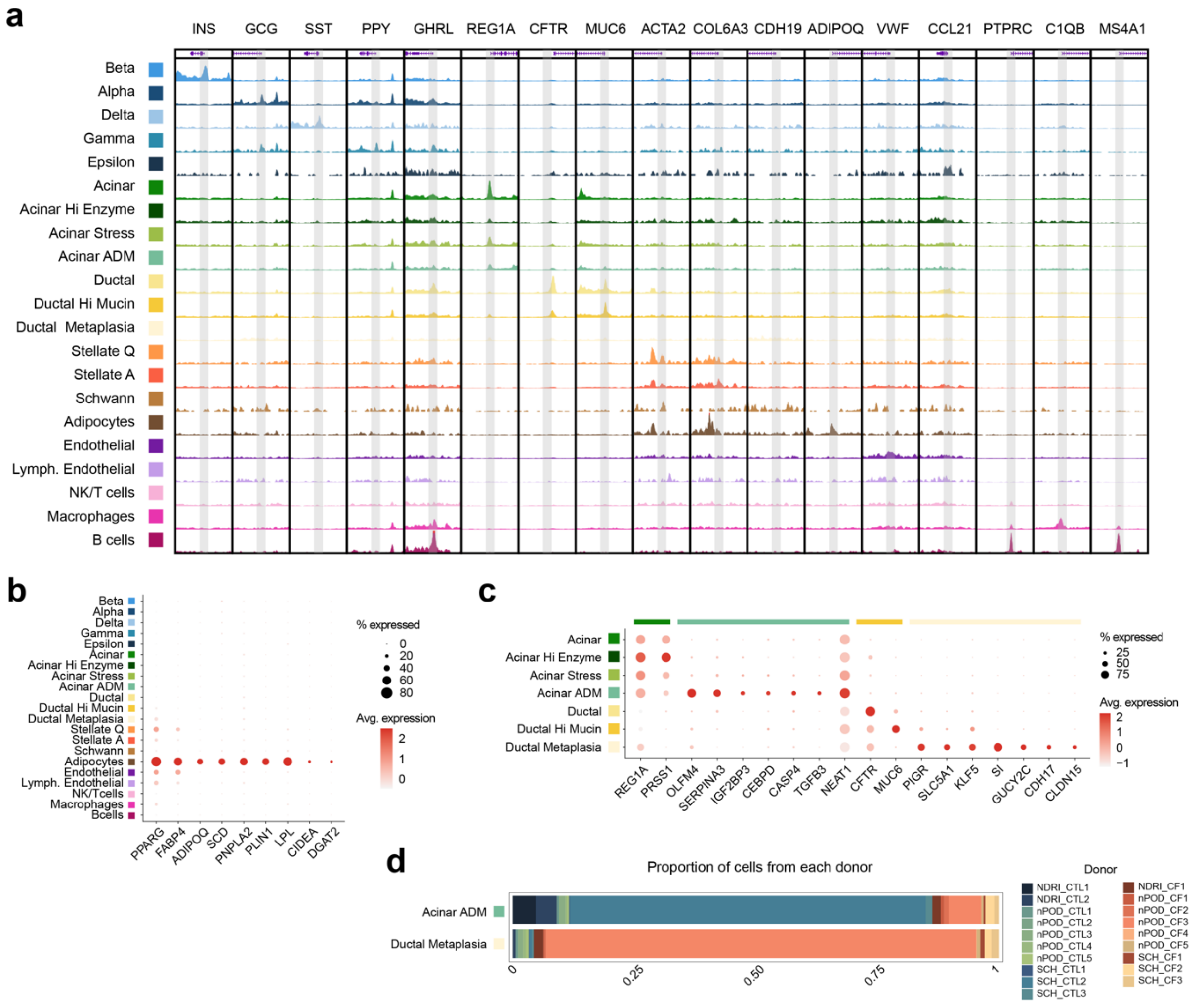


**Supplementary Figure 1. Chromatin and gene expression signatures of pancreatic cell types.** a) ATAC signal tracks for all cell type-marker genes. Each window contains a 6kb region centered around the marker gene transcription start site (TSS) and peak heights are standardized across all cell types for each region. b) All cell type gene expression of identified Adipocyte marker genes. The dot color represents the average expression in a cell type, and the dot size represents the percentage of cells within each cell type with non-zero expression. c) Exocrine cell type gene expression of markers of select cell types. The dot color represents the average expression in a cell type, and the dot size represents what percent of cells in the cell type have non-zero expression. The cell type each marker gene corresponds to is indicated by the colored bars at the top of the plot.

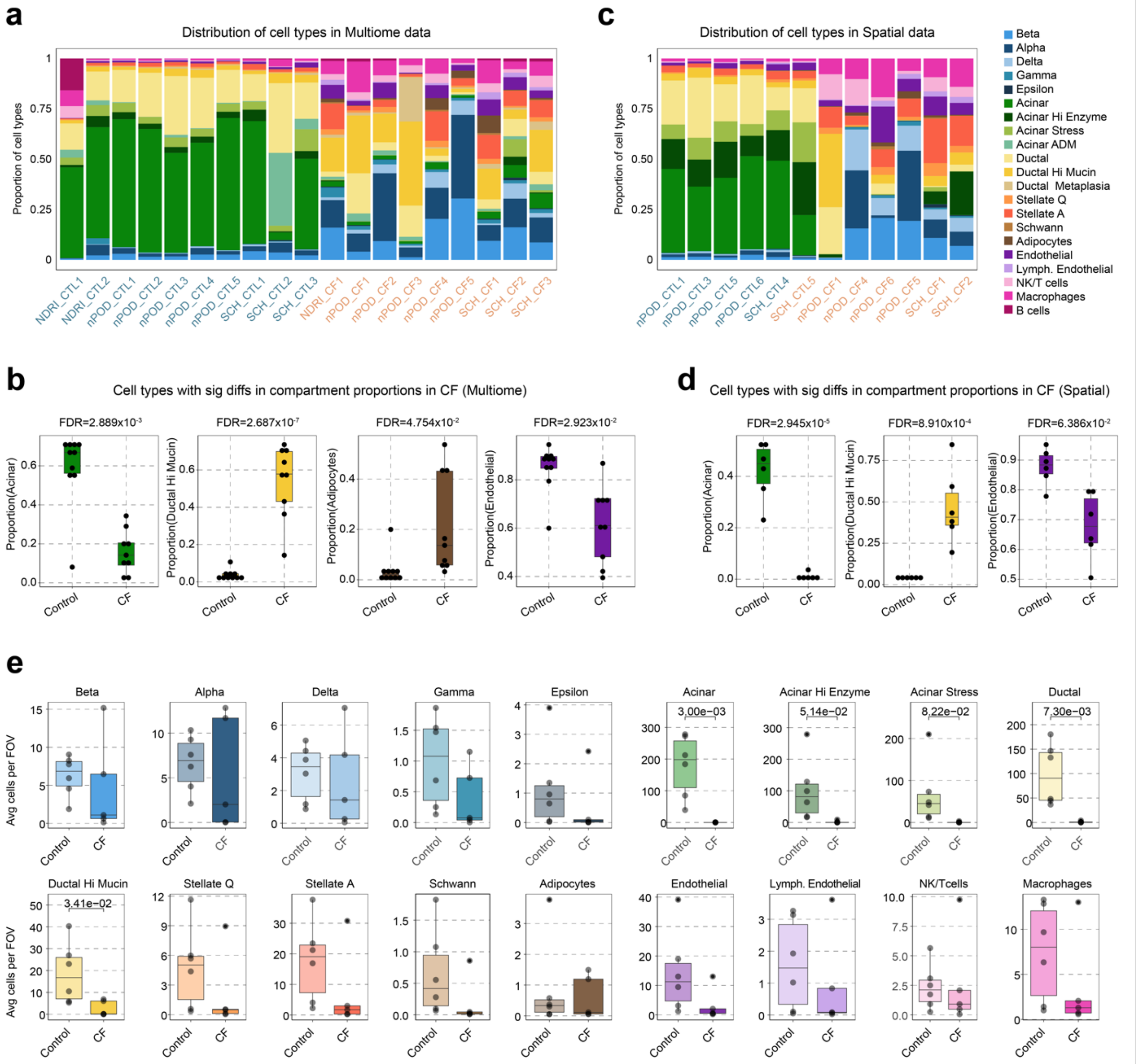

**Supplementary Figure 2. Cell type proportions changing with donor CF status.** a) Distribution of cell type proportions and sample covariate information across all study donors profiled via Multiome. b) All cell types with significant differences in scaled proportions between control and CF donors profiled via Multiome. Each boxplot represents the median (center line), first and third quartiles (bottom and top of box, respectively) and the limits of 1.5x interquartile range (vertical lines). Per-donor proportion values are marked as black dots. c) Distribution of cell type proportions and sample covariate information across all study donors profiled via CosMx. d) All cell types with significant differences in scaled proportions between control and CF donors profiled via CosMx. Per-donor proportion values are marked as black dots. e) The distribution of per donor average number of cells found in a CosMx spatial transcriptomics FOV region for all major cell types from Control and CF donors. Per donor counts are indicated as black dots. Each boxplot represents the median (center line), first and third quartiles (bottom and top of box, respectively) and the limits of 1.5x interquartile range (vertical lines).

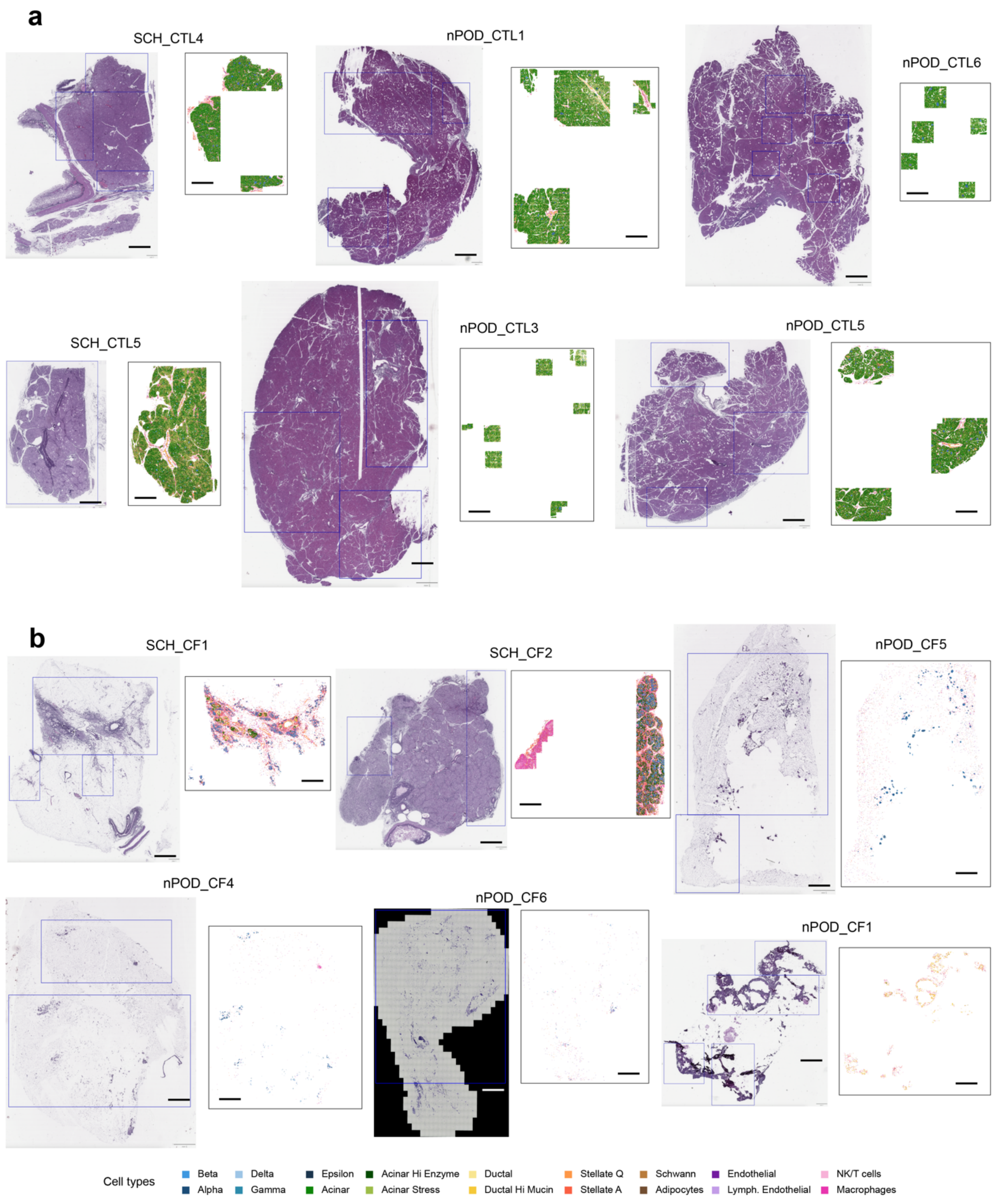


**Supplementary Figure 3. H&E images for all donor samples profiled with CosMx spatial transcriptomics.** a-b) Side by side images of H&E staining and spatial cell type coordinates for all control (a) and CF (b) donors. All scale bars represent 1mm and blue boxes on the H&E images correspond approximately to the tissue regions profiled with CosMx. All cell type colors are indicated by the key at the bottom.

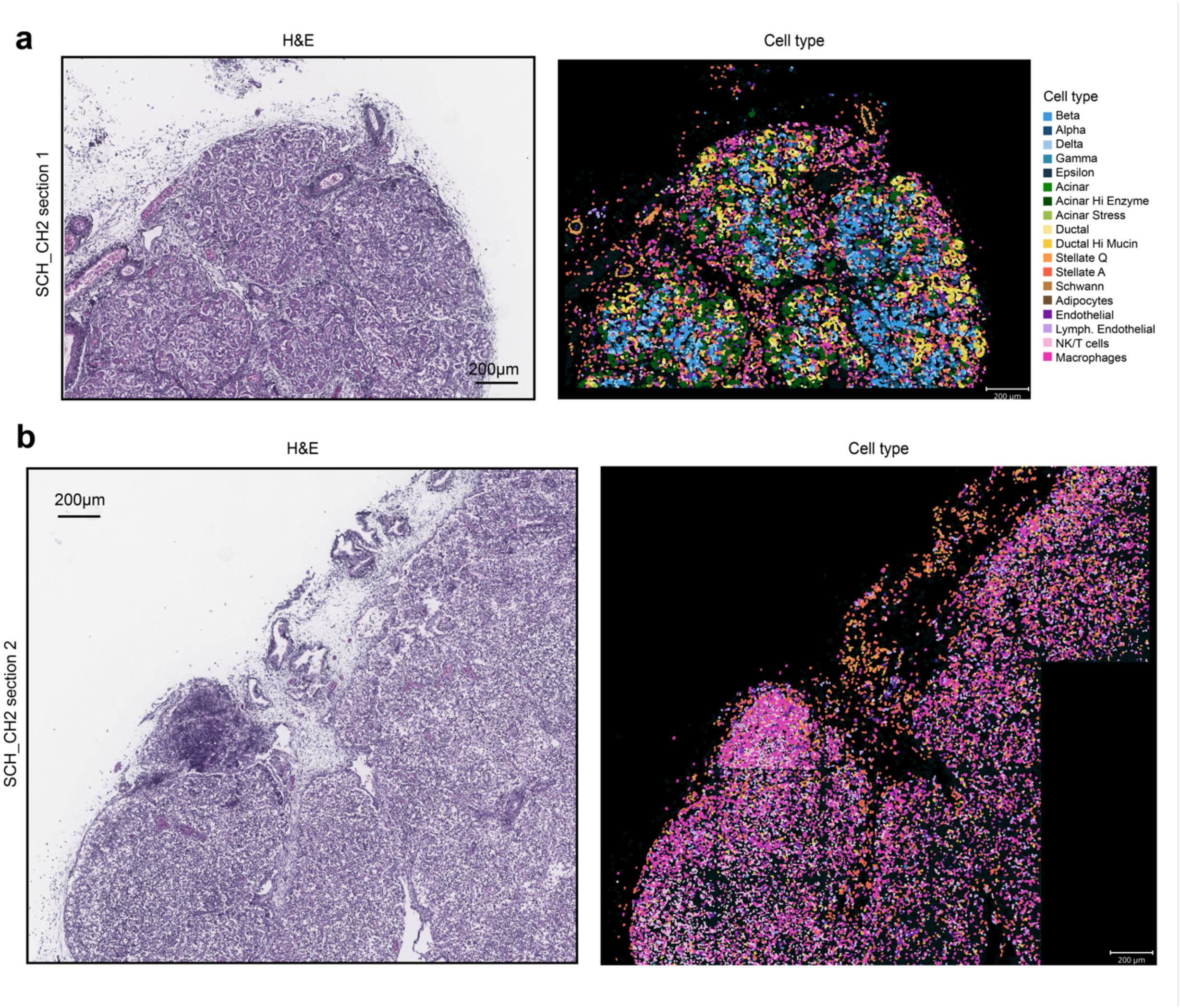


**Supplementary Figure 4. Tissue composition of pancreas from a young CF donor.** a-b) Representative images from two sections of tissue from donor SCH_CH2 with H&E staining (left column) and cell segmentation boundaries colored by imputed cell type assignments (right). All scale bars correspond to 200μm.

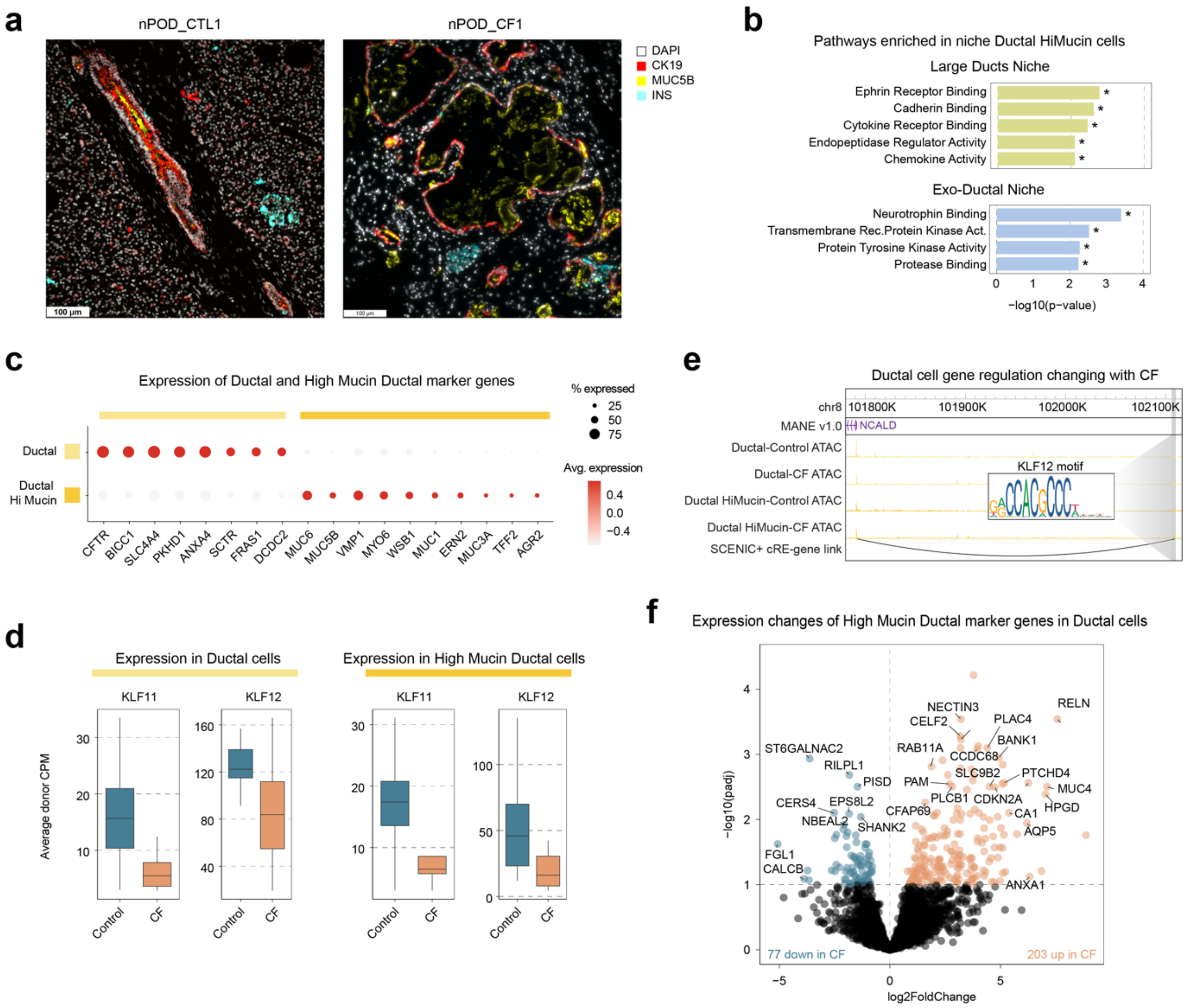


**Supplementary Figure 5. Expression of ductal cell marker genes in CF.** a) Select protein staining images from a control (left panel) and a CF donor (right panel), highlighting the differences in location and abundance of high-mucin ductal cells. b) Pathways enriched in genes upregulated in niche high-mucin ductal cells compared all other high-mucin ductal cells for two niches (top: Ducts, bottom: Exo-Ductal). c) Gene expression of ductal and high mucin ductal cell markers. The dot color represents the average expression in a cell type, and the dot size represents what percent of cells in the cell type have non-zero expression. The cell type each marker gene corresponds to is indicated by the colored bars at the top of the plot. d) Distribution of per-donor CPM-normalized gene expression of KLF TF family members in ductal and high mucin ductal cells, split by control and CF donors. e) Example TF-peak-gene trimer with increased activity in CF for high mucin ductal cells. f) Differential expression volcano plot composed of only high mucin ductal cell markers in ductal cells. All genes with differential expression passing

FDR<0.1 are colored and the vertical line demarks this cutoff. Genes with increased expression in CF are colored orange and genes with decreased expression in CF are colored blue.

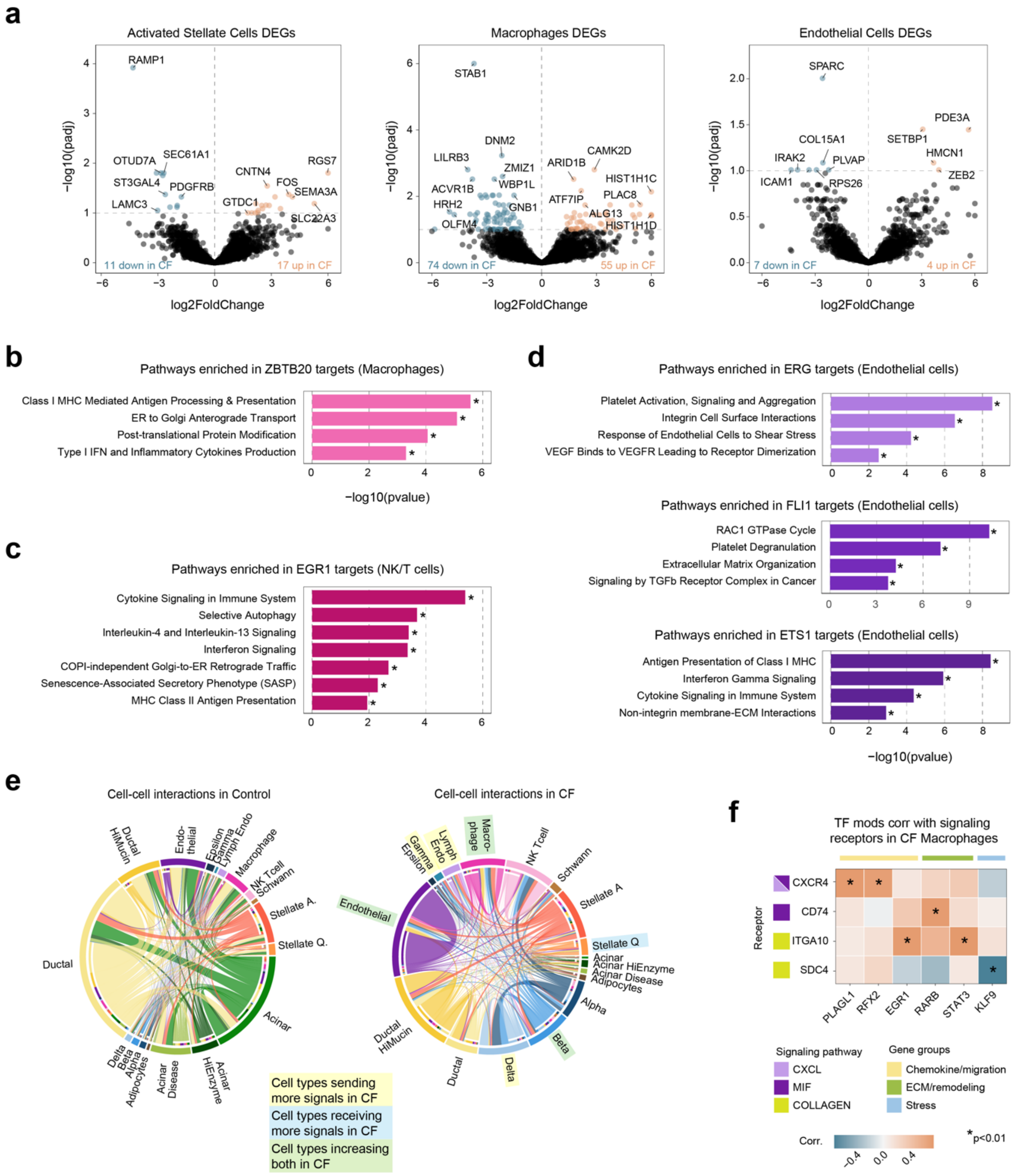


**Supplementary Figure 6. Differences in gene expression and signaling of inflammatory and structural pancreatic cell types in CF.** a) Volcano plots of genes with differential expression in CF donors for activated stellate cells (left), macrophages (middle), and endothelial cells (right). All genes with differential expression passing FDR<0.1 are colored and the vertical line demarks this cutoff. Genes with increased expression in CF are colored orange and genes with decreased expression in CF are colored blue. b-c) pathways enriched in the target genes of select TFs with changed expressions of target genes in endothelial cells (b) and immune cells (c) in CF. Significant enrichments (FDR<0.1) are marked with *. d) Summary of cell-cell signaling interactions between all cell types in control (left) and CF (right) pancreas. All chords originate from the sender cell type and connect to the receiver cell type, indicated by the smaller ring of colored bars. Cell types with nominally significantly altered (p<0.05) amounts of total signals sent or received are colored yellow and blue, respectively or green if both are true. e) Rank-based correlation of per-donor expression of signaling receptors with average per-donor SCENIC+ TF mod AUC values in CF macrophages. Nominally significant correlations (p<0.01) are marked with *.

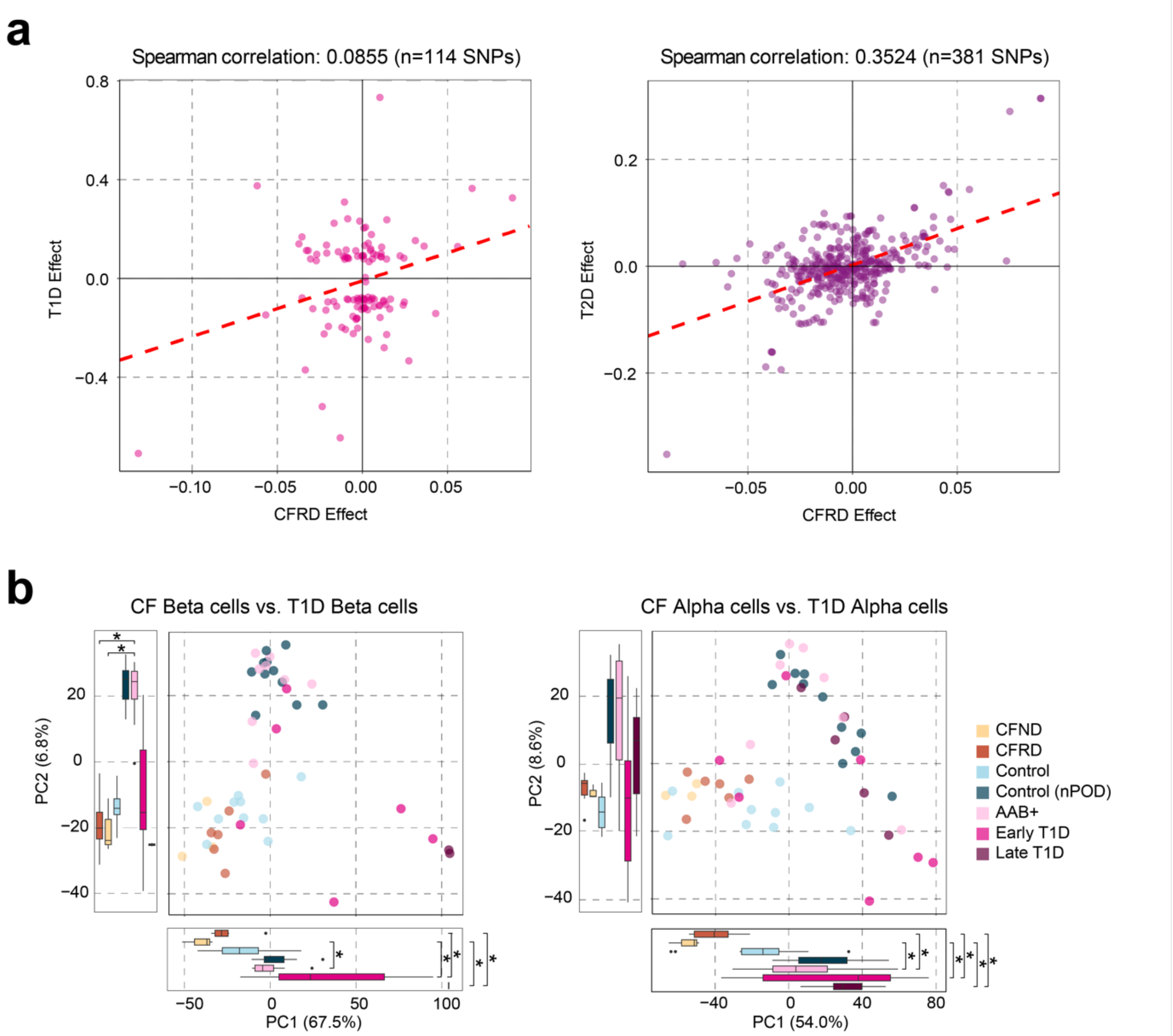


**Supplementary Figure 7. Comparison of CFRD risk and gene expression signatures to other forms of diabetes.** a) Direct comparison of effect sizes of SNPs significantly associated with risk of T1D (left) or T2D (right) with effect on CFRD risk. Each dot represents a T1D/T2D lead SNP also found in CFRD sumstats from an external publication. b) PCA comparison of donor pseudobulk RNA profiles with pancreatic autoantibodies (AAB+), early T1D, or late T1D from an external dataset. A type 2 ANOVA and then a Tukey's test were run to identify groups of donors with different mean PC embeddings. All $p<0.05$ differences between a CF group and a T1D group are annotated with *.

**Supplementary Tables and Data**

**Supplementary Table 1. Characteristics of donors profiled in this study**
**Supplementary Table 2. Marker genes of all identified pancreatic cell types**
**Supplementary Table 3. SCENIC+ TF modules enriched in specific cell types**
**Supplementary Table 4. Associations between cell type proportions and CF status**
**Supplementary Table 5. Genes associated with CF status**
**Supplementary Table 6. Pathways enriched in genes associated with CF status**
**Supplementary Table 7. TF modules enriched in CF-associated genes**
**Supplementary Table 8. Sequence motif accessibility associated with CF status**
**Supplementary Table 9. cRE accessibility associated with CF status**
**Supplementary Table 10. Genes and TF modules correlated with CF ductal cell trajectory analysis**
**Supplementary Table 11. Changes in cell type proximity to tissue features in CF pancreata**
**Supplementary Table 12. Pathways enriched in genes correlated with distance to tissue features in CF cell types**
**Supplementary Table 13. Cell-cell proximity changes in CF for pancreatic cell types**
**Supplementary Table 14. Signaling pathways with overall changes in CF**
**Supplementary Table 15. Cell types with changes in the total amount of signals sent or received in CF**
**Supplementary Table 16. Pairs of cell types with changes in signaling in CF**
**Supplementary Table 17. Cell type pairs with changes in specific signaling pathways in CF**
**Supplementary Table 18. Genes and TF modules correlated with signal receptor expression in Macrophages**
**Supplementary Table 19. Connectivity analysis comparisons of gene sets**
**Supplementary Table 20. Per-islet measurements different in CF pancreata**
**Supplementary Table 21. Per-islet measurements different based on CF islet environment**
**Supplementary Table 22. Pathways and TF modules correlated with signal receptor expression in Beta cells**
**Supplementary Table 23. Immunofluorescence antibodies**

**Supplementary Data 1. SCENIC+ eRegulons**
**Supplementary Data 2. Per-sample signaling pathway probabilities**
**Supplementary Data 3. Per-islet measurements**
